# MscM uses a novel gating mechanism for bacterial mechanosensitive channels

**DOI:** 10.64898/2026.03.24.713958

**Authors:** Giorgos Hiotis, Thomas Walz

## Abstract

The mechanosensitive channel of small conductance (MscS) is the founding member of the family of MscS-like channels, which share a structurally conserved core but feature additional structural elements that define their specific channel characteristics. Here, we characterize the structure and function of the *Escherichia coli* mechanosensitive channel of mini conductance (MscM), which features eight additional transmembrane (TM) helices and a large periplasmic domain. Our cryo-EM structures reveal that channel gating involves conformational changes in all domains of MscM. In particular, a cytoplasmic extension of TM7 couples the conformation of the TM domain to that of the cytoplasmic domain, resulting in gating of its lateral fenestrations, where ions enter the channel. Thus, different from all other MscS-like channels studied to date, channel gating in MscM is mediated by its cytoplasmic domain and not the TM domain, which senses changes in membrane tension and operates the cytoplasmic gates.

## INTRODUCTION

Mechanosensitive (MS) channels are membrane proteins that conduct ions upon mechanical activation^1–3^. In eukaryotic organisms, MS channels are responsible for a diverse set of functions, including roles in cell division, adhesion, somatosensation and hearing^1,2^. In bacteria, MS channels are activated during hypoosmotic shocks, serving as emergency release valves for efflux of solutes, preventing cell lysis^3,4^. Two types of MS channels have been discovered in bacteria: the MS channel of large conductance (MscL) and the MS channel of small conductance (MscS)^3,5^. Both channels are activated by directly sensing changes in membrane tension^6^. This observation gave rise to the “force-from-lipid” principle, which posits that changes in the transmembrane (TM) pressure profile can activate MS channels^6^.

Several structures of *E. coli* MscS (*Ec*MscS) have been determined by X-ray crystallography and cryogenic electron microscopy (cryo-EM), detailing its architecture and gating cycle^7–10^. *Ec*MscS assembles into a homoheptamer, with distinct TM and cytoplasmic domains^7–10^ (Fig. 1a). Each monomer contributes an N-terminal amphipathic helix and three TM helices. TM1 and TM2 form the tension-sensing paddle^7–10^. TM3 has a kink midway through the helix, dividing it into TM3a and TM3b. TM3a forms the ion-conducting pore, while TM3b connects to the cytoplasmic cage, which contains side openings, or fenestrations, through which ions enter the channel and which define the ion selectivity of the channel^7–10^. Upon activation, the TM1-2 paddles tilt and rotate, leading to opening of the TM pore and allowing ion conductance. Sustained exposure to membrane tension leads to a further increase in the tilt and a reverse rotation of the TM1-2 paddle, resulting in an inactivated state, in which the TM pore is closed^9^.

**Figure 1.**
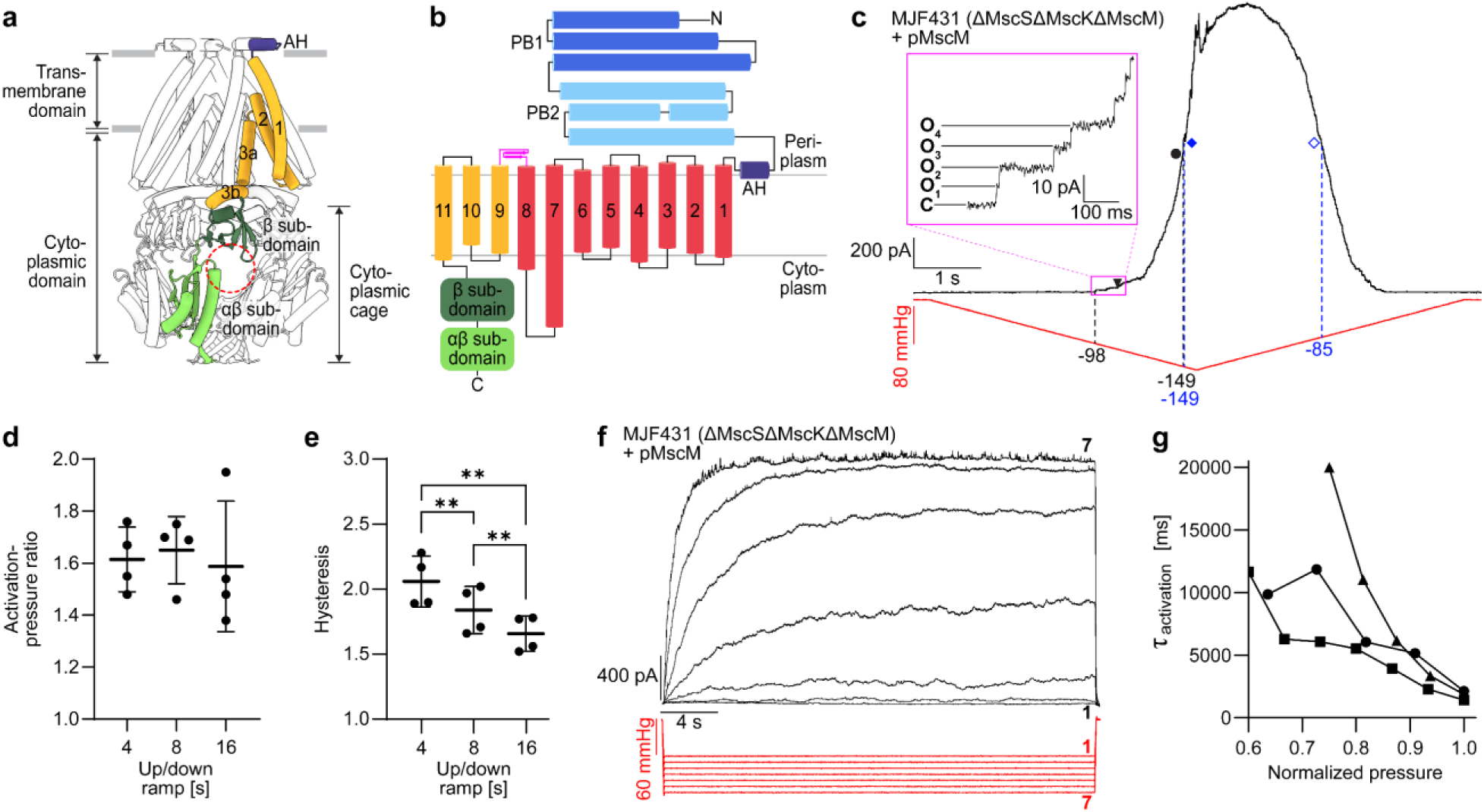
Topology and electrophysiological characterization of MscM in *E. coli* giant spheroplasts. **a** Structure of the archetypal *E. coli* MscS channel that consists of a cytoplasmic cage (light and dark green), TM helices 1-3 (yellow), and an amphipathic helix (AH) (purple). The dashed red circle indicates a cytoplasmic fenestration, through which ions enter the channel. **b** MscM consists of a large periplasmic domain that is organized into two distinct helical bundles, PB1 (dark blue) and PB2 (light blue), an amphipathic helix (AH) (purple), eleven TM helices, of which three are part of the MscS-like core structure (orange) and eight are additions in MscM (red), and a cytoplasmic domain that is composed of the β subdomain (dark green) and the αβ subdomain (light green). **c** Representative patch-clamp recording (n = 4) from a giant spheroplast generated from *E. coli* strain MJF431 (which does not express MscS, MscK and MscM) overexpressing MscM in response to a symmetric 8-s-long pressure ramp from 0 mmHg to -160 mmHg and back to 0 mmHg. Currents were recorded in the inside-out excised patch configuration with a pipette potential of +30 mV. Pressure applied to the patch membrane and the recorded current are shown in red and black, respectively. Black triangle: first opening of MscM; black circle: first opening of MscL; filled blue rhombus: mid-point of MscM; empty blue rhombus: mid-point deactivation of MscM; dashed black lines: pressures at which the first MscM channel and the first MscL channel open; dashed blue lines: pressures at which the mid-point activation and mid-point deactivation of MscM occur. The inset shows a zoomed-in view of the MscM currents in the region indicated by the purple box. C: current level when all MscM channels are closed; O_1_-O_4_: current levels when one to four MscM channels are open. **d** Quantitative analysis shows no significant changes in the activation-pressure ratio of MscM in response to ramps of different speeds. **e** Quantitative analysis shows significant changes in the hysteresis ratio of MscM in response to ramps of different speeds. Statistical significance was determined with a paired t-test (n = 4, ** *p* ≤ 0.001). **f** Representative patch-clamp recording (n = 3) from a giant spheroplast generated from *E. coli* strain MJF431 overexpressing MscM in response to 30-s-long pulses with the pressure increasing from -100 mmHg to -160 mmHg in -10 mmHg intervals. MscM activates slowly and shows no discernible desensitization/inactivation. **g** Analysis of the activation-time constant τ_activation_ of MscM in response to the maximum normalized pressure applied within each of three patches from recordings shown in panel d. MscM activation becomes faster with increasing pressure.

MscS is the archetypal member of the superfamily of MscS-like channels that all share a similar core structure with three TM helices and a cytoplasmic cage but show substantial diversity in additional structural elements and functional characteristics^5,11^. In addition to MscS, *E. coli* expresses five MscS-like channels^5,11^. These channels differ in their number of TM helices. YnaI and YbdG have five TM helices, while YbiO, MscM (or YjeP) and MscK (or KefA) have eleven TM helices and also feature extensive periplasmic domains^5^. Previous studies showed that MscS responds best to fast osmotic changes, while other MscS-like channels respond better to slow osmotic changes^12–14^.

MscM and MscK, the largest MscS-like channels in *E. coli* (∼1100 amino acids; a.a.), feature eleven TM helices per monomer and large periplasmic domains (∼450 a.a.) that are organized into two distinct helical bundles, PB1 and PB2 (Fig. 1b). Previous studies demonstrated that MscK is a potassium-dependent MS channel, the first example of an MscS-like channel having requirements for activation in addition to membrane tension^15^. Recent cryo-EM structures of MscK revealed the architecture and gating of this channel^16^. In the resting state, the periplasmic domain assembles into a ring-like structure, and the TM domain is significantly curved. Channel gating is accompanied by a dilation of the periplasmic ring, a flattening and expansion of the TM domain, and an increase in the pore radius from 2.6 Å to 9.3 Å.

MscM appears to be very similar to MscK. The sequence and predicted topology of MscM very closely resemble those of MscK. The two channels have 29% sequence identity and 48% similarity, with particularly high sequence homology in TM11 and the β subdomain (75% identity, 89% similarity). Despite these similarities, their electrophysiological characteristics differ. Previous patch-clamp electrophysiology recordings from *E. coli* giant spheroplasts showed that the conductance of MscM is smaller than that of MscK (∼0.3 nS *versus* ∼0.9 nS)^11,15^. Additionally, the pressure sensitivity of MscM is almost identical to that of MscS, while MscK is the most pressure-sensitive MS channel in *E. coli*^11^. Furthermore, while gating of MscK has been shown to depend on periplasmic potassium^15^, no such ion dependency has yet been established for MscM.

Here, we report cryo-EM structures of MscM in the closed and open conformations. In the closed conformation, the membrane-proximal periplasmic PB2 domains assemble into a ring and the TM domain is curved. Upon opening, the PB2 ring disassembles and the TM domain expands and becomes completely flat, increasing the pore radius from 1.8 Å to 10.2 Å. Notably, our structures show that a cytoplasmic extension of TM7 structurally and functionally couples the TM domain to the cytoplasmic domain. In particular, the lateral fenestrations in the cytoplasmic cage are closed in the resting state, but flattening of the TM domain is transmitted by TM7 to the cytoplasmic cage, which undergoes a conformational change that results in the opening of the fenestrations. Thus, unique to MscM, it is not the TM pore but the fenestrations in the cytoplasmic cage that function as the channel gates.

## RESULTS

### Functional characterization of MscM

We used patch-clamp electrophysiology to analyze the functional characteristics of MscM expressed in *E. coli* strain MJF431^4^. This strain, in which MscS-like channels MscS, MscK and MscM are knocked out, still expresses MscL, which can thus be used for comparison with MscM. We prepared giant spheroplasts^14,16,17^ and applied symmetric pressure ramps (4 s up / 4 s down) to excised inside-out patches to a maximum pressure sufficient to see activation of both MscM and MscL (110 - 160 mmHg, depending on the patch) (Fig. 1c). The activation of multiple MscM channels at lower pressure, seen as distinct steps in the current curve over time (Fig. 1c, inset), allowed us to calculate its single-channel conductance as 258 ± 102 pS (mean ± standard deviation; n = 4), which is similar to the previously reported value for the conductance of MscM^11^. We then determined the activation-pressure ratio for MscM, defined as the ratio between the pressures at which the first MscL channel and the first MscM channel open, P_MscL:MscM_, as 1.62 ± 0.12. This activation-pressure ratio is very similar to a previously determined value for MscM^11^ (1.64 ± 0.07), as well as for MscS^18^ (1.62 ± 0.03), suggesting that MscM and MscS open at a comparable membrane tension. Our recordings also revealed that MscM displays strong gating hysteresis, which we quantified as the ratio of the mid-point opening pressure to the mid-point closing pressure. The hysteresis ratio we measured for MscM is 2.06 ± 0.20, which is considerably higher than that of MscS^19^ (∼1.25). The use of slower pressure ramps (8 s up / 8 s down and 16 s up / 16 s down) revealed that the activation pressure is not meaningfully affected by the ramp speed (Fig. 1d), whereas the gating hysteresis decreased with slower ramp speeds (2.06 ± 0.20, n = 4 for the 4 s / 4 s ramp; 1.84 ± 0.18, n = 4 for the 8 s / 8 s ramp; and 1.66 ± 0.14, n = 4 for the 16 s / 16 s ramp) (Fig. 1e).

To evaluate whether MscM desensitizes/inactivates, we applied to the same excised inside-out patches a series of 30-s-long pressure pulses with increasing strength (Fig. 1f). The recordings revealed that MscM activates slowly over the duration of the pulses, with MscM channels continuing to open until the pressure is released. No desensitization/inactivation was apparent in any of the recordings. To quantify the activation kinetics of MscM and to investigate whether different pressure levels affect it, we fit exponential curves to each pulse in our recordings and determined the opening time constant τ_open_, the time it takes to reach 63.2% of maximum current. This analysis revealed that τ_open_ decreases, i.e., MscM activation becomes faster, with increasing pressures (Fig. 1g). These findings show that the activation kinetics of MscM differ from those of MscS, which activates rapidly (with peak MscS currents occurring within a few tenths of milliseconds) but then desensitizes/inactivates as the pressure persists over several seconds^20^.

### Structure of MscM in the closed state

We expressed MscM in *Pichia pastoris* and purified the recombinant protein in a buffer containing glycol-diosgenin (GDN) and 150 mM NaCl (Fig. S1a,b). Cryo-EM analysis yielded a density map at an overall resolution of 6.7 Å resolution (Fig. S1c-f). The map only revealed interpretable density for the cytoplasmic domain and TMs 7-11 (Fig. S1g). The density for TMs 1-6 and the periplasmic domain did not resolve individual helices. Furthermore, the density representing the periplasmic domain was too small to include the entire domain, suggesting that it represented at most the second helical bundle, PB2 (Fig. 1b). Because the map did not show density for PB1, we tested whether removing PB1 (MscM ΔPB1) (Fig. S2a) would affect the electrophysiological properties of the channel. Our recordings revealed that removing PB1 had no statistically significant effect on the single-channel current, activation pressure and hysteresis of the channel, nor did it alter its slow activation kinetics and lack of desensitization/inactivation (Fig. S2b-f) and so we removed it from expression constructs for further structural studies.

Cryo-EM analysis of MscM ΔPB1 yielded a map at an improved overall resolution of 4.3 Å that resolved the cytoplasmic domain and all the helices of the TM and PB2 domains (Fig. 2a and Fig. S3a-d), allowing us to build a backbone model for the entire channel except PB1 (Fig. 2b and Fig. S3i). 3D classification without alignment and local refinement focusing on the MscS-like core structure of MscM, namely TMs 9-11 and the cytoplasmic domain, improved the overall resolution of the core structure to 3.4 Å (Fig. S3e-h). However, only the cytoplasmic domain and TM11 were sufficiently well resolved to build an atomic model (Fig. S3i). We also locally refined the core structure of full-length MscM and used it to build an atomic model for the cytoplasmic domain and TM11 (Fig. S1h-l). Alignment of the cytoplasmic domain and TM11 from full-length MscM and MscM ΔPB1 revealed that the two structures are nearly identical with a root-mean-square deviation (RMSD) between backbone atoms of 0.294 Å (Fig. S3j), supporting the notion that truncation of PB1 does not affect the overall structure of MscM.

**Figure 2.**
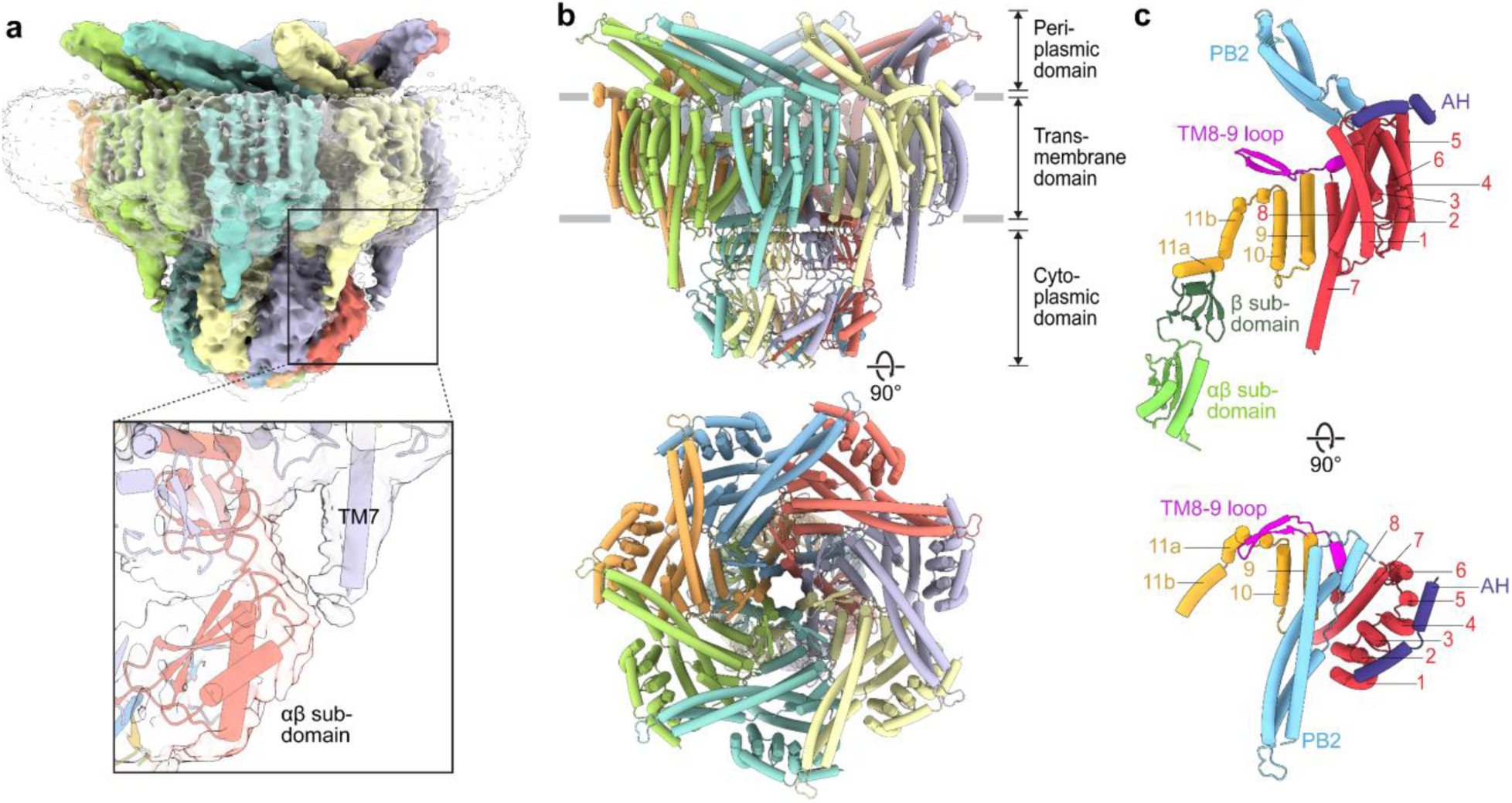
Cryo-EM structure of MscM ΔPB1 in a closed conformation. **a** Cryo-EM map of MscM ΔPB1 at 4.3-Å resolution colored by subunit. Density representing the detergent micelle is shown as transparent surface. The inset shows a zoomed-in view of the cryo-EM map (transparent surface) and model of MscM (purple and red cartoon) of the interaction between the cytoplasmic extension of TM7 and the αβ subdomain. **b** Views parallel (top) and perpendicular (bottom) to the membrane of MscΜ ΔPB1 shown in cartoon representation and colored by subunit. **c** Views parallel (top) and perpendicular (bottom) to the membrane of an MscM ΔPB1 colored and labeled according to the topology shown in Fig. 1b. For clarity, the cytoplasmic domain is not shown in the view perpendicular to the membrane.

The model of MscM ΔPB1 reveals the organization of the periplasmic, TM and cytoplasmic domains (Fig. 2b,c). The seven PB2s, which assemble into a ring structure, connect to TM1 through an amphipathic helix, AH, that is localized at the periphery of TMs 1-6. The extensive classification that was necessary to resolve PB2 and TMs 1-6 as well as their lower resolution compared to the rest of the structure, suggest that they are dynamic. TMs 1-6 form a helical bundle whose outside surface faces the membrane and inside surface interfaces with TM7 (Fig. 2b,c). TM7 has a long cytoplasmic extension and connects to TM8 through a loop, whose density was too weak to allow modeling. However, the density connects to the αβ subdomain of the cytoplasmic domain of the neighboring monomer, suggesting a domain-swapped direct interaction between them (Fig. 2a, inset). TM8 connects to TM9 through an extended periplasmic loop that contains a pair of short antiparallel β-strands that interact with each other. While density for this loop is weak, it is close to the three TM helices of the core structure, potentially linking pore-lining TM11 to TM7 (Fig. 2c). Like in all MscS-like channels, pore-lining TM11 contains a kink that divides it into TM11a and TM11b. TM11a forms the ion-conducting pore of the channel, while TM11b connects to the cytoplasmic domain, which is organized into a cage-like structure featuring lateral fenestrations that allow ions to enter the cytoplasmic domain. The overall TM domain is curved, as is evident from both 2D-class averages (Figs S1c and S3a) and the 3D-density map (Fig. 2a). This bent conformation must deform the surrounding membrane, which is energetically unfavorable. Such membrane deformation has also been observed for other MS channels, like MscK^16^ and PIEZO1^21,22^, indicating that MscM may be activated in a similar manner as those channels.

Analysis of the pore radius using HOLE^23^ revealed an extended constriction region formed by residues Trp902, Val909 and Phe913 (Fig. 3a, left and middle panels). These residues arrange into ring structures at the periplasmic end, middle and cytoplasmic end of the pore, where it narrows to radii of 1.8 Å, 3.1 Å and 2.0 Å, respectively, suggesting that the pore is in a closed conformation. We therefore will refer to MscM ΔPB1 structure as representing the closed conformation.

**Figure 3.**
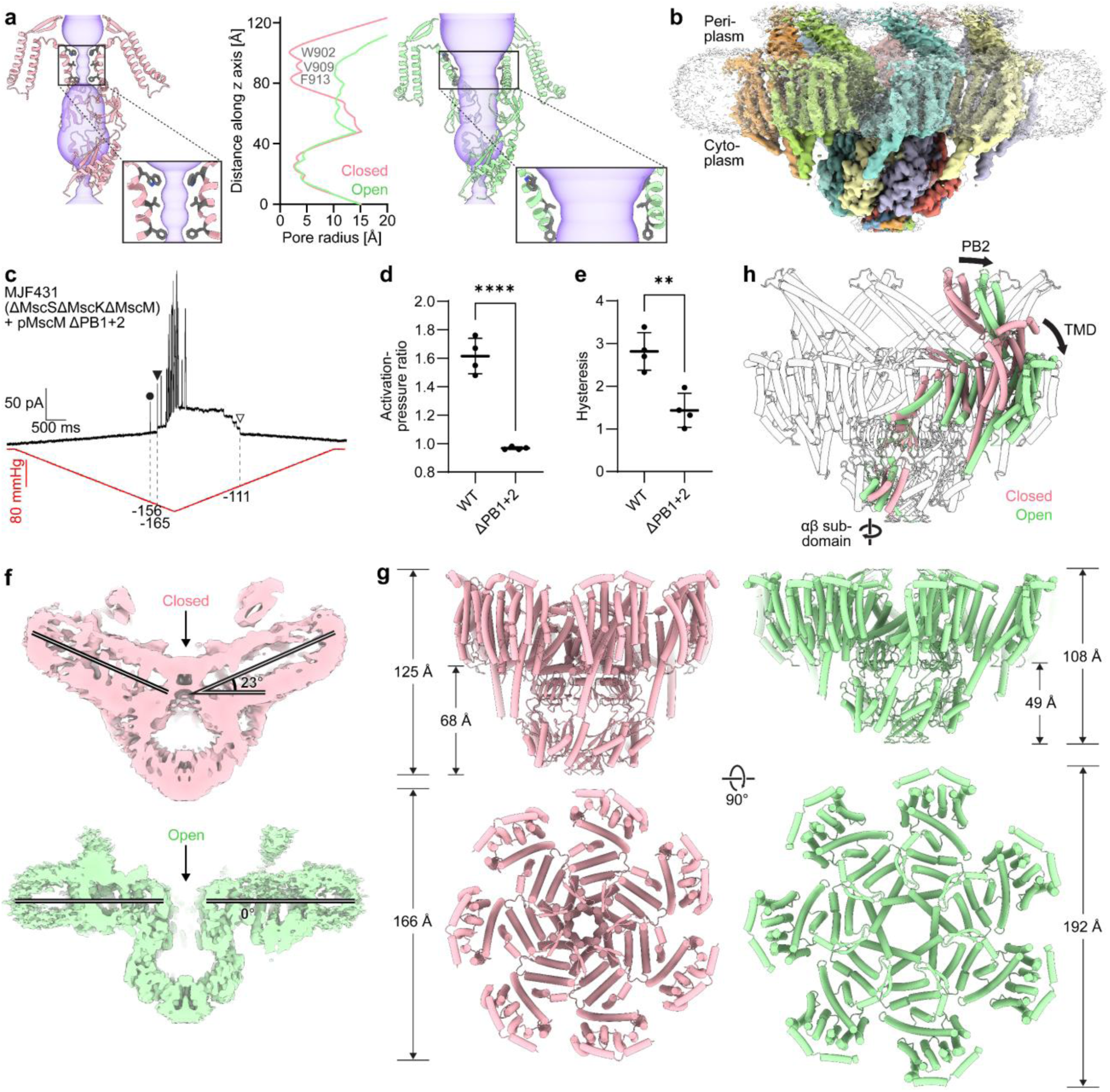
Cryo-EM structure of WT MscM in an open conformation and structural changes associated with channel gating. **a** Analysis of the pore radius of MscM in the closed conformation (left; pink ribbons) and the open conformation (right; green ribbons) using HOLE^23^. Residues Trp902, Val909 and Phe913, whose side chains form the narrowest constriction points in the ion-conduction pathway (transparent purple surfaces), are shown as gray sticks. The middle panel shows a plot of the pore radius of MscM along the z axis in the closed (pink line) and open conformation (green line). Opening of MscM increases the narrowest pore radius from 1.8 Å to 10.2 Å. **b** Cryo-EM composite map of WT MscM in an open conformation colored by subunit. Density representing the detergent micelle is shown as transparent surface. The periplasmic bundles PB2 are no longer in contact with each other, and the TM domain is flat. **c** Representative recording (n = 4) from a giant spheroplast generated *from E. coli* strain MJF431 (which does not express MscS, MscK and MscM) overexpressing MscM ΔPB1+2 in response to a symmetric 8-s-long pressure ramp from 0 mmHg to -190 mmHg and back to 0 mmHg. Currents were recorded in the inside-out excised patch configuration with a pipette potential of +30 mV. Pressure applied to the patch membrane and the recorded current are shown in red and black, respectively. Filled black circle: first opening of MscL; filled black triangle: first opening of MscM; empty black triangle: last opening of MscM; dashed black lines: pressures at which the first and last MscM and the first MscL channels open. **d** Quantitative analysis shows a significant reduction in the activation-pressure ratio of MscM ΔPB1+2 compared to that of WT MscM. Statistical significance was determined with an unpaired t-test (n_WT_ = 4, n_ΔPB1+2_ = 4, **** *p* ≤ 0.0001). **e** Quantitative analysis shows a significant reduction in the hysteresis ratio of MscM ΔPB1+2 compared to that of WT MscM. Statistical significance was determined with an unpaired t-test (n_WT_ = 4, n_ΔPB1+2_ = 4, ** *p* ≤ 0.01). **f** Cut-away views parallel to the membrane of the cryo-EM maps of MscM in the closed (top panel, pink surface) and open conformation (bottom panel, green surface), showing that the TM domain of MscM in the open conformation is completely flat. The arrows point to the TM pore. **g** Dimensions of the TM and cytoplasmic domains of MscM in the closed (left panel, pink) and open conformation (right panel, green). Views are parallel (top) and perpendicular to the membrane (bottom). When MscM opens, the αβ subdomain partially drops into the TM domain, which flattens and expands, thus substantially increasing the footprint of MscM. For clarity, the periplasmic domain is not shown in both views, and also the cytoplasmic domain is not shown in the view perpendicular to the membrane. **h** Overlay of the structure of MscM in the closed (one subunit shown in pink) and open conformation (one subunit shown in green and the other six subunits in white). The arrows indicate conformational changes associated with the gating of MscM.

### Structure of MscM in the open conformation

To determine the structure of MscM in the open conformation, we first introduced the G912S mutation in pore-forming TM11, which corresponds to the gain-of-function mutation G924S in MscK that was used to determine its structure in the open conformation^16^. However, cryo-EM analysis showed that this mutant remained in the closed conformation (Fig. S4). Therefore, we explored different buffer conditions as a potential means to open MscM. When we purified MscM in buffer containing 150 mM KCl, cryo-EM 2D-class averages revealed the channel in two distinct conformations, one in which the overall TM domain is curved, as seen in the structure of MscM in the closed conformation, and one in which the TM domain is essentially flat (Fig. S5a). Image processing, including extensive 3D classifications, yielded a map of MscM with a flat TM domain at an overall resolution of 2.8 Å (Figs S5a-d and S6). While the map only resolved TMs 7-11 and the cytoplasmic domain, it did reveal a large TM pore compared to the closed TM pore in MscM ΔPB1 (Fig. 3a, right panel), establishing that a flat TM domain indicates MscM in the open conformation. Analysis of the pore radius using HOLE revealed a dilated pore with radii of 14.1 Å, 10.4 Å and 10.2 Å at the three constriction points formed by residues Trp902, Val909 and Phe913 (Fig. 3a, middle panel).

By employing symmetry expansion and focused classification, we were able to resolve all the TM helices in a single monomer of the channel at an overall resolution of 3.2 Å (Fig. S5e-h), but the resolution of TMs 1-6 was only sufficient to model the protein backbone (Fig. S6). We also performed iterative rounds of heterogeneous refinements to remove particles with no density representing the periplasmic domain, which eventually yielded a map at an overall resolution of 3.3 Å that showed clear density for the periplasmic domain, albeit with low local resolution (Fig. S5i-l). While the map showed density for PB2, individual helices were not resolved, and so we used rigid-body docking to place the AlphaFold model of PB2 into the density (Fig. S6). The final composite map shows that the PB2 domains of MscM in the open conformation are no longer in contact with each other (Fig. 3b), providing a reason for the increased flexibility of PB2 and TMs 1-6 compared to those of MscM in the closed conformation.

We used electrophysiology to probe the role of the periplasmic ring formed by the PB2s. Recordings obtained with a mutant MscM lacking both periplasmic helical bundles, MscM ΔPB1+2, revealed that membrane patches contained only few MscM channels (∼1 - 5 channels per patch) (Fig. 3c). The pressure required to open MscM ΔPB1+2 was much higher than that needed for wild-type (WT) MscM and close to that required for MscL (Fig. 3d). Occasionally, the first MscM channel opened even after the first MscL channel opened. Furthermore, truncation of the entire periplasmic domain also significantly reduced gating hysteresis of the channel compared to WT MscM (Fig. 3e). While the low number of channels in the patches prevented us from characterizing the activation kinetics of MscM ΔPB1+2, it is clear that deletion of the periplasmic PB2 ring substantially decreases the mechanosensitivity of MscM and reduces the delay in channel closing seen for WT MscM.

The transition from the closed to the open conformation involves conformational changes in all three domains of MscM (Fig. 3f-h and Movie S1). Channel opening is accompanied by a flattening and expansion of the TM domain, decreasing the midplane bending angle from ∼23° to ∼0° (Fig. 3f) and increasing the diameter and area of its in-plane footprint from 166 Å to 192 Å and from 21,642 Å^2^ to 28,953 Å^2^, respectively (Fig. 3g). While TMs 1-10 primarily undergo a rigid-body movement, TM11, which forms the ion-conducting pore of the channel, loses the kink that is observed in the closed conformation and becomes continuous (Fig. 3g). These changes move the PB2s further away from the channel axis and further apart from each other than in the closed conformation (Fig. 3h), so that they can no longer form a ring structure (Fig. 3b) and increasing the mobility of the individual PB2s. Most notably and in contrast to all other studied MscS-like channels, the cytoplasmic domain also undergoes a conformational change, with the αβ subdomain rotating clockwise with respect to the β subdomain (Fig. 3h and Movie S2). This new conformation of the cytoplasmic domain is more similar to the cytoplasmic domain conformation of MscS and all other MscS-like channels (see below).

We also collected a cryo-EM dataset of MscM ΔPB1 purified in buffer containing 150 mM KCl. Intriguingly, this dataset also showed part of the MscM population in the open conformation (Fig. S7), providing further evidence that a high potassium concentration can induce MscM to adopt the open conformation and that channel opening does not require PB1.

### TM7 coordinates the conformations of the TM and cytoplasmic domains

The different conformations of the cytoplasmic domain seen in the structures of MscM in the closed and open states are stabilized by several unique backbone and side-chain interactions between the β and the αβ subdomains (Fig. S8a,b). Furthermore, the cryo-EM maps of MscM in the closed and open conformations suggest that the cytoplasmic TM7 extension and the TM7-8 loop interact with the αβ subdomain of the cytoplasmic domain. Even though the density map did not allow us to model these interactions, we hypothesized that the connection of the TM7 extension and TM7-8 loop with the αβ subdomain might couple the conformation of the TM domain to that of the cytoplasmic domain. An AlphaFold-Multimer^24^ prediction of MscM placed the TM7-8 loop in close proximity to the αβ subdomain, as seen in our cryo-EM maps, and predicted interactions of TM7-8 loop residues His754, Ser757 and Glu759 with residues lining the cytoplasmic fenestrations (Fig. S8c). However, the missing density for the TM7-8 loop in all our maps indicates that these interactions, if real, would not be strong enough to stabilize the conformation of the loop.

The AlphaFold-Multimer prediction shows PB2 forming a ring structure, like in our map of MscM in the closed conformation, and PB1 forming a second ring that covers the periplasmic surface of the PB2 ring (Fig. S8d). A comparison of the AlphaFold prediction with our structure of MscM in the closed conformation shows the TM domain and PB2 in similar positions (Fig. S8e). However, while pore-lining TM11 is essentially the same, suggesting that the AlphaFold model represents a closed conformation, the cytoplasmic αβ subdomains are not aligned. Alignment of the core structure of the AlphaFold model with our structures of MscM in the closed and open conformations revealed that AlphaFold did not predict the novel closed conformation we observed for the cytoplasmic domain of MscM (Fig. S8f).

Since we could not identify the exact residues that mediate the interaction between the TM and the cytoplasmic domain, we deleted the parts of the TM7 extension and the TM7-8 loop most likely to interact with the αβ subdomain (amino acids 741-759) (Fig. S8g) and analyzed this ΜscM ΔPB1ΔΤΜ7e mutant by cryo-EM. The resulting map at an overall resolution of 5.1 Å resolved the periplasmic and cytoplasmic domains as well as all TM helices (Fig. 4a and Fig. S9a-d), allowing us to build backbone models for all domains (Figs 4b and S9i). As expected, the map showed no density for the truncated cytoplasmic extension of TM7. Local refinement of the MscM core structure yielded a map for this region at a resolution of 3.5 Å that allowed us to build an atomic model for TMs 9-11 and the cytoplasmic domain (Fig. S9e-h). Removing the interaction of the TM domain with the cytoplasmic domain resulted in two notable differences.

**Figure 4.**
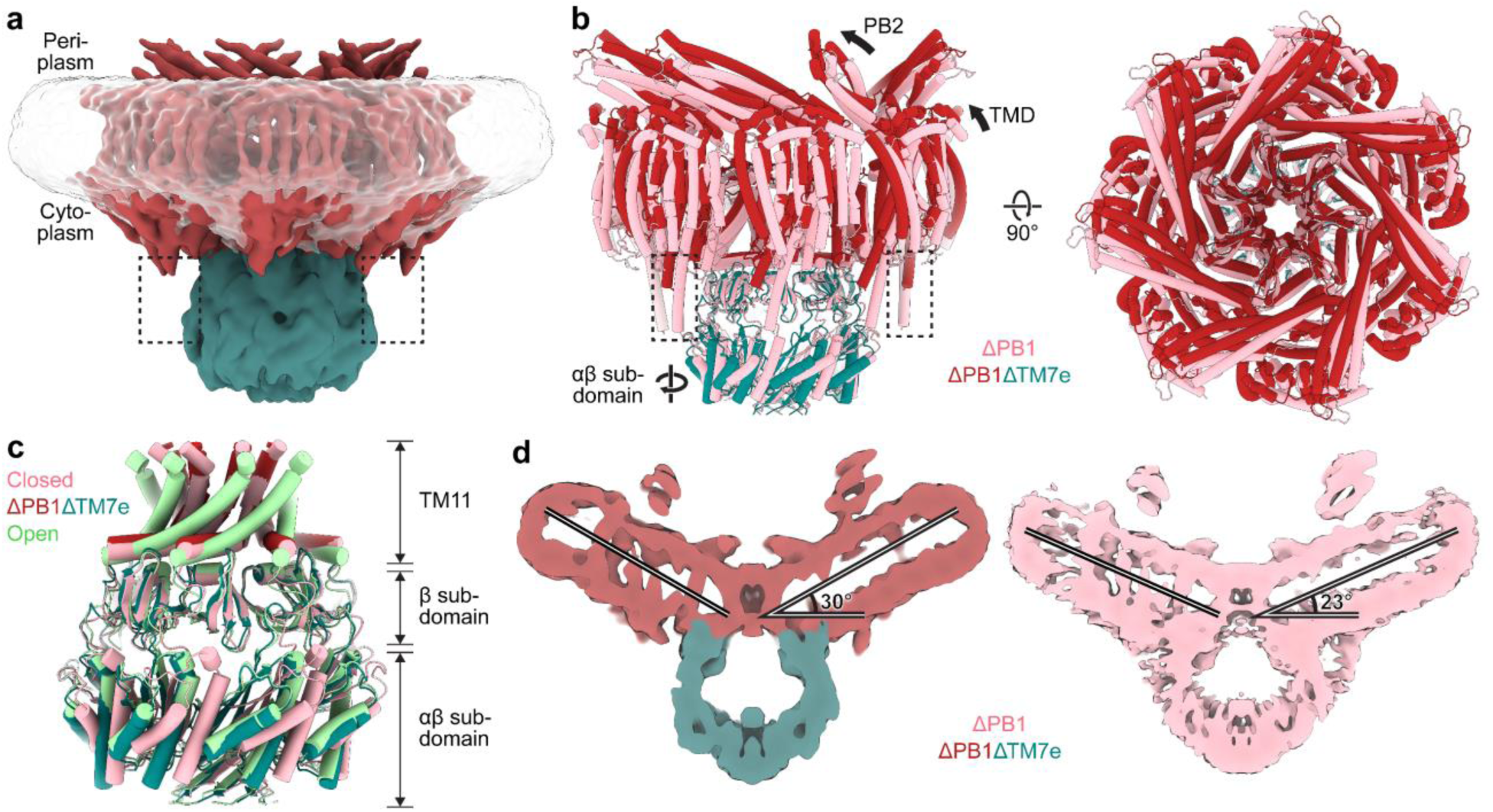
The role of TM7 in the conformational coupling of the TM and cytoplasmic domains. **a** Cryo-EM structure of MscM ΔPB1ΔTM7e colored by the conformation of each domain. The periplasmic and TM domains are in the closed conformation (dark red), while the cytoplasmic domain adopts the open conformation (teal). The truncated TM7 does not interact with the αβ subdomain, as indicated by the dotted boxes. Density representing the detergent micelle is shown as transparent surface. **b** Views parallel (left) and perpendicular (right) to the membrane of the structural alignment of MscM ΔPB1 in the closed conformation (pink) and MscM ΔPB1ΔTM7e with the closed periplasmic and TM domains (dark red) and open cytoplasmic domain (teal), showing that truncation of TM7 (dotted boxes) leads to conformational changes in all domains of MscM. **c** Structural alignment of TM11 and the cytoplasmic domain of MscM ΔPB1ΔTM7e (red and teal) with MscM in the closed (pink) and open (green) conformations, showing that the truncation of TM7 leads to the cytoplasmic domain adopting the same conformation as seen in the open conformation, while TM11 remains in the closed conformation. **d** Cut-away views parallel to the membrane of the cryo-EM maps of MscM ΔPB1ΔTM7e (left; periplasmic and TM domains in red and cytoplasmic domain in teal) and MscM ΔPB1 in the closed conformation (right; pink), showing that truncation of TM7 increases the curvature of the TM domain.

As hypothesized, the cytoplasmic domain adopts the conformation seen for MscM in the open conformation (Fig. 4c and Movie S3), but, unexpectedly, the TM domain has a more pronounced curvature, with a midplane bending angle of ∼30°, larger than the 23° seen for MscM in the closed conformation (Fig. 4d and Movie S3). This result confirms that TM7 indeed couples the conformations of the TM and cytoplasmic domains, demonstrating that extended TM7 is responsible not only for the unconventional conformation of the cytoplasmic domain of MscM in the closed conformation but also affects the curvature of the TM domain in the closed conformation.

### The fenestrations in the cytoplasmic domain constitute a novel second gate

We used molecular-dynamics (MD) simulations to investigate whether the MscM fenestrations have a different ion permeability in the closed and open conformations. We used the core structures of MscM ΔPB1, which has closed fenestrations and a closed TM pore (Fig. 5a), and ΜscM ΔPB1ΔΤΜ7e, which has open fenestrations and a closed TM pore (Fig. 5b), and for comparison, the structure of MscS in the closed conformation^9^ (Fig. 5c) (PDB: 6VYK). We placed the channels in a solution containing 1 M KCl and monitored the increase in the number of ions in the cytoplasmic domain over time (Fig. 5d). Over the course of the simulations, the total number of ions in the cytoplasmic domains of MscS and MscM with open fenestrations rapidly increased and plateaued over the last 500 ns at 53.4 ± 0.1 and 35.5 ± 4.6 (average ± standard deviation; n = 3) total ions, respectively. In contrast, only 12.5 ± 4.2 ions entered the cytoplasmic domain of MscM with closed fenestrations. The cumulative count of ions permeating the fenestrations (into and out of the cytoplasmic domains) correlates well with the size of the respective fenestrations, showing a very fast increase in the cumulative count for MscS, a slower increase for MscM in the open conformation, and almost no increase for MscM in the closed conformation (Fig. 5e). These trends remain the same when looking separately at ions entering and exiting the cytoplasmic domain through the fenestrations (Fig. S10a,b) and are further illustrated by simulation snapshots from the mid-point and last frames of the simulations (Fig. S10c).

**Figure 5.**
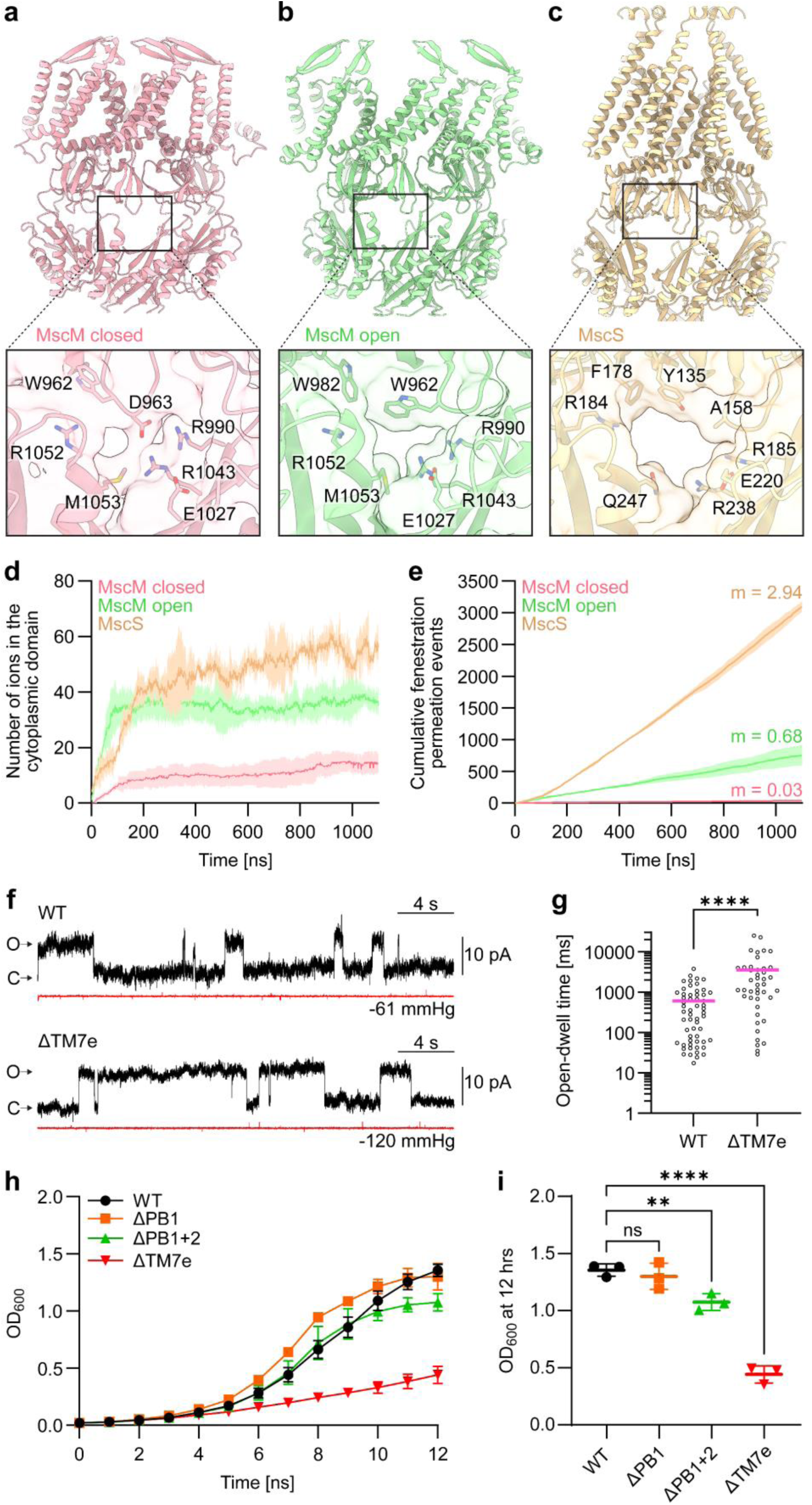
The fenestrations of MscM constitute a novel second gate. a-c. Structures of MscM in the closed (a) (pink ribbons) and open conformation (b) (green ribbons), and MscS in the closed conformation^9^ (c) (yellow ribbons; PBD: 6vyk). Insets show a zoomed-in view of a fenestration. Maps are shown as transparent surfaces to show the size of the openings and residues lining the fenestrations are represented as sticks. Opening of MscM increases the size of the fenestrations, but these are still smaller than those of MscS. **d** Plot of the number of ions that occupy the cytoplasmic domains of MscM with closed (red) and open (green) fenestrations and MscS (yellow) for the duration of the simulations. The lines represent the mean value and the shaded areas the standard deviation (n = 3). **e** Plot of the cumulative penetration events (both entry and exit) through the fenestrations of MscM with closed (red) and open (green) fenestrations and MscS for the duration of the simulations. The lines represent the mean and the shaded areas the standard deviation. The numbers indicate the slope for each line obtained from linear regression models. **f** Representative recordings from giant spheroplasts generated from *E. coli* MJF431 strain overexpressing either WT MscM (top panel) or MscM ΔTM7e (bottom panel) in response to 30-s-long pressure pulses sufficient to see single-channel opening events. C and O indicate the current levels resulting from closed and one open MscM channel. The open-dwell time of the MscM ΔΤΜ7e mutant appears longer than that of WT MscM. **g** Quantitative analysis shows that the open-dwell time of MscM ΔΤΜ7e is significantly higher than that of WT MscM. Each data point represents the open-dwell time of a single event pooled across multiple recordings from multiple patches. Statistical significance was calculated using a Mann-Whitney U-test (**** *p* ≤ 0.0001). **h** Growth curves of *E. coli* MJF641 (which does not express any MS channels) cultured in M9 minimal medium and expressing WT MscM (black line and circles) or truncation mutants ΔPB1 (orange line and squares), ΔPB1+2 (green line and upward triangles) or ΔTM7e (red line and downward triangles). The plot shows the OD_600_ of the cultures over the course of 12 hours. Expression of ΔΤΜ7e leads to a pronounced growth delay. **i** Analysis of the OD_600_ of all constructs at the 12-hours time point. Statistical significance was determined with a one-way ANOVA test (** *p* ≤ 0.01, **** *p* ≤ 0.0001). Expression of MscM ΔPB1 does not significantly delay growth of *E. coli.* Expression of MscM ΔPB1+2 causes a modest yet significant growth delay, while expression of MscM ΔTM7e leads to a substantial growth delay.

The MD results not only confirm that the fenestrations in the cytoplasmic domain of MscM constitute a novel second gate but also demonstrate that the MscM fenestrations, even in the open conformation, are much less ion-conductive than those of MscS. This finding explains why MscM has a three-fold lower ion conductance compared to MscS, despite its open TM pore being wider (radius ∼10 Å) than that of MscS^8^ (radius ∼6 Å). To study the contribution of the cytoplasmic gate to the channel properties of MscM, we used electrophysiology to characterize the MscM ΔTM7e deletion mutant and found no appreciable differences in activation pressure, gating hysteresis and activation kinetics compared to those of WT MscM (Fig. S11). However, the recordings suggested that MscM ΔTM7e has substantially longer open-dwell times (Fig. 5f). To better characterize the open-dwell times, we recorded several current curves at a pressure that caused only single-channel openings. Quantification revealed that the open-dwell time increased significantly from 611 ± 764 ms for WT MscM to 3,560 ± 5,437 ms for MscM ΔTM7e (average ± standard deviation, n_WT_ = 57 from 3 patches, n_ΔΤΜ7e_ = 43 from 5 patches; p < 0.0001, Mann-Whitney U test) (Fig. 5g). These data thus suggest a role for the TM7-mediated interaction of the TM domain with the cytoplasmic domain in limiting the open-dwell time of MscM.

To investigate this possibility, we used an approach previously used to determine the impact of gain-of-function mutations in MscK on the growth of *E. coli*^25^. We transformed *E. coli* strain MJF612, in which all MS channels are knocked out^26^, with pTRC99a encoding WT MscM or MscM ΔPB1, ΔPB1+2 and ΔTM7e. The cells were grown in minimal medium, protein expression was induced after four hours, and growth was monitored every hour for an additional eight hours (Fig. 5h,i). Upon induction of protein expression, cells carrying the pTRC99a vector with MscM ΔTM7e grew substantially slower compared to cells carrying either pTRC99a with WT MscM or MscM ΔPB1. Less substantial but still significantly slower growth was seen for cells carrying pTRC99a with MscM ΔPB1+2. While we did not see spontaneous gating activity with the MscM ΔTM7e mutant in electrophysiology, which could have explained the observed growth defect, these results show that in a physiological context, the lack of connection between the TM and cytoplasmic domains and the resulting open conformation of the cytoplasmic gate leads to a deleterious dysregulation of MscM.

### The gating mechanisms of MscM and MscK differ

MscM is highly homologous to MscK, and structural comparison shows the expected similarity in the overall architecture of the two channels in the closed conformation (Fig. S12a-c). Both channels have periplasmic domains of comparable size (MscK: 53.7 kDa, MscM: 49.9 kDa), with the membrane-distal periplasmic helical bundle PB1 not resolved in the cryo-EM maps and the membrane-proximal periplasmic helical bundle PB2 assembling into a ring. The eleven TM helices per monomer have comparable arrangements in MscK and MscM, but a key difference is that TM7 in MscM has a long cytoplasmic extension. Because of its shorter TM7 and the larger in-plane bending angle of the TM domain in the closed conformation, the TM7 of MscK does not interact with the αβ subdomain of the cytoplasmic cage (Fig. S12b). Structural alignment of the core structures of MscM and MscK reveals that their TM11 helices share a similar arrangement, but their cytoplasmic domains adopt different conformations (Fig. S12d). The cytoplasmic domain conformation of MscM differs from that of MscK and all other structurally characterized MscS-like channels due to the rotation of the αβ subdomain relative to the β subdomain that is stabilized by the extension of TM7 and the TM7-8 loop.

The conformational changes associated with opening of MscM and MscK show several key differences (Fig. S12e-g and Movie S4). In MscK, the periplasmic ring formed by the PB2s persists upon opening, whereas the PB2s of MscM dissociate. The TM domains of both MscM and MscK expand and flatten as the channels transition into the open state, but while the TM domain of MscM almost completely flattens, the TM domain of MscK retains an in-plane bending angle of ∼22°, resulting in a smaller increase in its in-plane footprint (3,695 Å^2^) compared to that of MscM (7,311 Å^2^). Given that TM11 is highly conserved between MscM and MscK, their similar pore radii in the closed (MscM: 1.8 Å, MscK: 2.6 Å) and open conformations (MscM: 10.2 Å, MscK: 9.3 Å) are expected. In both channels, a conserved tryptophane residue, Trp902 in MscM and Trp914 in MscK, form a hydrophobic lock at the periplasmic side of the pore. The greatest differences associated with gating occur in the cytoplasmic domains of the two channels. As a result of the almost complete flattening of the TM domain in MscM, its cytoplasmic domain partially drops into the TM domain, and the flattening of the TM domain is transmitted through extended TM7 to the αβ subdomain, which rotates with respect to the β subdomain, opening the previously closed fenestrations (Movie S2). As a result, the conformation of the cytoplasmic domain of MscM in the open state becomes the same as that of MscK (Fig. S12h).

We also compared the structure of MscM ΔPB1ΔΤΜ7e to that of MscK, since deletion of the TM7 extension and the TM7-8 loop led to a higher degree of curvature in the TM domain compared to MscM without this deletion (Fig. 4d). An overlay shows that the TM domain of MscM ΔPB1ΔΤΜ7e and its curvature are much more similar to those of McsK in the closed conformation (Fig. S12i,j) than full-length MscM in the closed conformation (Fig. S12c). These comparisons establish the TM7 extension and the TM7-8 loop as the central elements that distinguish the structure and function of MscM from those of MscK.

## DISCUSSION

MscS-like channels are characterized by an MscS-like core and feature additional structural elements. *E. coli* expresses MscS and five MscS-like channels, raising the question what the physiological roles of all these channels are and how the additional structural elements modify their functional characteristics. YnaI is possibly the best characterized MscS-like channel. It features two additional TM helices, which make it much less mechanosensitive^11,14,27^. A recent electrophysiological characterization revealed that YnaI has slow gating kinetics, which makes it much better suited to respond to slow changes in membrane tension than MscS, which reacts best to fast changes in membrane tension^14^. Furthermore, unlike MscS, YnaI has pronounced gating hysteresis and does not desensitize/inactivate, so that it remains open for a long time after initial activation, allowing for continued ion conduction long after MscS and MscL have already closed again^14^. Much less is understood for the other MscS-like channels.

MscM has eight additional TM helices per subunit and a large periplasmic domain. Electrophysiological characterization showed that MscM opens at a similar membrane tension as MscS but differs from MscS in all other aspects. It has slow gating kinetics, pronounced gating hysteresis, and does not desensitize/inactivate, making it more similar to *E. coli* YnaI^14^ and the plant MscS-like channels FLYC1^28^ and AtMSL10^29,30^. To understand the functional role of the additional structural elements in conferring functional characteristics, we deleted them individually and analyzed the effect by electrophysiology.

We first studied the membrane-distal periplasmic bundle PB1, which, according to AlphaFold, would form a ring that is nestled into the ring formed by the membrane-proximal periplasmic bundle PB2. Despite the predicted extensive interactions with the PB2 ring, PB1 was not visible in any of our MscM structures nor in any of the previously published structures of MscK, which also has a PB1 bundle^16^. Deletion of PB1 in MscM had no effect on any of the functional characteristics we analyzed. These findings suggest that PB1 may not be arranged as predicted by AlphaFold and may not be essential for the core functions of MscM. We note, however, that a search for MscM and MscK genes in the Uniprot database^31^ found homologs only in Gram-negative bacteria, which are characterized by an outer cell membrane. It is thus possible that PB1 can only exert its effects on MscM function in the context of a well-defined periplasmic space and an outer cell membrane, which are missing in the spheroplasts used for patch-clamp electrophysiology.

Removal of the entire periplasmic domain, PB1 and PB2, significantly reduced the mechanosensitivity of MscM as well as its gating hysteresis. Since the channel characteristics did not change when only PB1 was deleted, these effects must be caused by the loss of PB2. PB2 connects to the TM1-6 bundle through amphipathic helix AH, raising the possibility that it helps maintain the curvature of the TM domain, a feature that makes MS channels sensitive to membrane tension^22,32^. In support of this notion, YbiO, which also has eleven TM helices per subunit but has a much smaller periplasmic domain, is much less mechanosensitive than MscM and MscK^11^. Removal of PB2 from MscM would potentially allow the TM domain to reduce its curvature without opening the TM pore, thus explaining why the deletion mutant is less mechanosensitive than WT MscM. Concerning the reduced gating hysteresis observed for the deletion mutant, PB2 forms a ring in the closed conformation, which dissociates when the channel opens. The lack of interactions between the PB2s and the resulting increased mobility of the periplasmic domain in the open conformation would make it difficult for the PB2s to reassemble into a ring as the channel closes. The asymmetric pathways, a simple one for ring disassembly and a more complex one for ring reassembly, could explain how PB2 contributes to the gating hysteresis of MscM and how deletion of PB2 would diminish hysteresis.

We also removed all the additional TM helices of MscM, TMs 1-8, but we could not detect expression of this mutant, suggesting that the MscS-like core in MscM has evolved too far away from MscS to be capable of still forming a stable, membrane-inserted channel.

The structure of MscM in the closed conformation shows the expected similarity in topology and architecture with MscK, but comparison revealed two salient differences, which are the cytoplasmic extension of TM7 in MscM that contacts the cytoplasmic αβ subdomain and the lesser curvature of its TM domain. Understanding the functional relevance of these differences required knowing the structure of MscM in the open conformation. The first attempt to open the channel, by introducing the same mutation in the pore-lining helix that allowed the structure of MscK to be determined in the open conformation, failed to induce an open conformation in MscM, providing a first indication that, despite the similarity between the two channels, MscM may be gated differently from MscK. Similarly, while potassium is required but not sufficient for MscK to gate^15^, potassium was sufficient to obtain the structure of detergent-solubilized MscM in the open conformation, further supporting the notion that the two channels gate differently.

For MscK to gate, potassium must be present on the periplasmic side, identifying the periplasmic domain as the potassium sensor. Here, we show that potassium also opens MscM lacking membrane-distal helical bundle PB1, thus establishing that membrane-proximal helical bundle PB2 is the potassium-sensitive element. In MscM in the open conformation, the PB2s no longer form a ring, indicating that potassium binding to PB2 destabilizes the PB2 ring and that the PB2 ring may play a role in stabilizing the closed conformation. The observation that the PB2 ring does not dissociate in MscK in the open conformation may potentially explain why potassium alone is sufficient to gate MscM but not MscK in detergent, and why MscK lacks the strong gating hysteresis present in MscM^16^.

Studies on MscL^33,34^ and PIEZO1^22,32^ established that the mechanosensitivity of a channel is proportional to the increase in the in-plane membrane area and the decrease in membrane curvature upon channel opening. Our structures reveal that MscM has a larger increase in membrane footprint when it transitions from the closed to the open conformation than MscK, ΔA_MscM_ = 7,311 Å^2^ *versus* ΔA_MscK_ = 3,695 Å^2^, as well as a larger decrease in induced membrane curvature, Δθ_MscM_ = 23° *versus* Δθ_MscK_ = 14° (note that in the transition from the closed to the open state, the TM domain of MscM flattens completely, the curvature changes from 23° to 0°, whereas the TM domain of MscK remains curved, the curvature changes from 36° to 22°). Based on these observations, MscM should be more mechanosensitive than MscK, but the opposite is true (P_MscL_:P_MscK_ = 1.92^16^ *versus* P_MscL_:P_MscM_ = 1.64). This discrepancy implies that the closed conformation of MscM must be more stable than that of MscK. The main structural difference between the two channels is the cytoplasmic extension of TM7, which connects the TM domain to the cytoplasmic domain and may explain why the closed conformation of MscM is more stable than that of MscK. Intriguingly, in the MscM ΔTM7e mutant, the curvature of the TM domain is higher in the closed conformation and more similar to that of MscK, ∼30°. Assuming the open conformation of the MscM ΔTM7e mutant is the same as WT MscM, it would undergo a more substantial increase in membrane area expansion and decrease in TM domain curvature, which should render MscM ΔTM7e more mechanosensitive than WT MscM. However, its mechanosensitivity remains unchanged, suggesting that additional factors affect the sensitivity of MS channels to membrane tension.

The most unique feature revealed by our MscM structures is the conformational change in the cytoplasmic domain that has so far not been seen for any other MscS-like channel. To date, the function of the cytoplasmic domain was thought to be exclusively to define the ion selectivity of the channel^35^ and to form a rigid scaffold that functions as a stable anchor against which the TM helices can move. Our MscM structures now show that the β subdomain suffices as a rigid scaffold against which not only the TM domain can move but also the αβ subdomain. This notion is consistent with other MscS-like channels that entirely lack the αβ subdomain, such as the MscS-like channels from *Nematocida displodere*^36^ and *Trypanosoma cruzi*^37^. The αβ subdomain is, however, essential to form the lateral fenestrations and thus for the ion selectivity of the channels. Our MscM structures show that the fenestrations can function not only as passive ion filters but also as gates, expanding the list of roles the cytoplasmic domain can play.

Our electrophysiological characterization of MscM has several implications for the physiological role of MscM in hypoosmotic shock protection. The slow gating kinetics would render MscM more suitable to respond to slow osmotic changes, which allows more MscM channels to open over time, thus providing increasing protection as membrane tension persists. Intriguingly, MscK requires periplasmic potassium to open, and MscM seems even more sensitive to potassium than MscK. But where does the potassium come from? It is possible that the periplasmic potassium needed to open MscK and MscM was released from the bacterium itself by other potassium-independent MS channels that opened more quickly in response to a hypoosmotic shock, primarily MscS and MscL. However, the periplasmic potassium concentration would likely only increase locally and quite transiently, so that only MscK and MscM channels very close to those other channels would be activated, rendering this hypothesis unlikely. Alternatively, the potassium concentration in the environment may increase more globally if it were released by bacteria in the surrounding environment that lysed in response to hypoosmotic shock. In either case, the potassium dependence of MscK and MscM may suggest that these channels are a second line of defense after other MS channels had already been activated. Once activated, however, due to its pronounced gating hysteresis and its lack of desensitization/inactivation, MscM remains open for a long time and could reduce membrane tension much more than other MscS-like channels like MscS and YnaI, allowing membrane tension to return to a much safer level.

## METHODS

### Cloning

For structural characterization, the DNA sequence of *E. coli* MscM with a C-terminal 3C-cleavable eGFP-10xHis tag was optimized for *Pichia pastoris* codon usage and synthesized into the pUC57 vector (Bio Basic, Inc). The DNA fragment encoding tagged MscM was cloned into a pPICZb vector for expression in *P. pastoris*.

For functional characterization, the DNA sequence for *E. coli* MscM was cloned directly from *E. coli* genomic DNA into a pTRC99a vector.

All single-point and truncation mutants were performed with the Q5 Site-Directed Mutagenesis Kit (New England Biolabs), using primers designed with NEBaseChanger (https://nebasechanger.neb.com).

### Protein expression and purification

All MscM constructs were expressed in *P. pastoris* SMD1163H (Invitrogen). 30 ml of overnight cultures grown in yeast extract peptone dextrose (YEPD) medium were used to inoculate 1-l cultures in buffered minimal glycerol (BMG) medium and incubated at 30°C until OD_600_ reached ∼15-20. The cultures were centrifuged at 3,000xg for 10 minutes at room temperature, and the cells were resuspended in buffered minimal methanol (BMM) medium, containing 0.5% (v/v) methanol. The cultures were incubated for 48 hours at 24°C and methanol was replenished 24 hours after initial induction. The cells were harvested by centrifugation at 3,000xg for 10 minutes at 4°C. Cell pellets were flash frozen in liquid nitrogen and stored at -80°C until further use.

The purifications of WT MscM, MscM G912S and MscM ΔPB1 in NaCl buffer were performed as follows. A cell pellet (∼40 g) was resuspended in 160 ml of Lysis Buffer containing 30 mM Tris-HCl, pH 7.4, 150 mM NaCl, 5 mM MgCl_2_, 10% (v/v) glycerol, 0.05 mg/ml DNase I (Worthington Biochemical) and a protease inhibitor cocktail tablet (Sigma-Aldrich). The cells were lysed by passing them three times through a microfluidizer (Avestin EmulsiFlex-C3), operating at a pressure of ∼20,000 PSI. The lysate was clarified by centrifugation at 4,000xg for 20 minutes at 4°C, and the supernatant was centrifuged at 100,000xg for 1 hour at 4°C to isolate the membranes. The pellet was resuspended in 45 ml of Resuspension Buffer containing 30 mM Tris-HCl, pH 7.4, 150 mM NaCl and 10% glycerol. Following Dounce homogenization, 5 ml of 10% (w/v) GDN (Anatrace) solution was added to the membrane suspension to a final concentration of 1%, and membranes were solubilized for 2 hours at 4°C with rotation. The sample was centrifuged at 40,000xg for 30 minutes at 4°C to remove insoluble material, and the supernatant was incubated with 4 ml of GFP-nanobody resin for 1 hour at 4°C with rotation. The resin was transferred into a gravity column, washed with 10 column volumes (CV) of Buffer A containing 30 mM Tris-HCl, pH 7.4, 150 mM NaCl, 0.02% GDN, supplemented with 10% glycerol. The column was capped, supplemented with 2 ml of Buffer A and 500 μl of 3C protease solution (∼3 mg/ml in 50 mM Tris pH 8, 150 mM NaCl and 1 mM DTT), and incubated for 2 hours at 4°C. The protein was eluted with 15 ml of Buffer A supplemented with 10% glycerol and concentrated to ∼0.5 ml using an Amicon Ultra concentrator with a 100-kDa molecular-weight cutoff by centrifugation at 4,000xg and 4°C. The protein was further purified by size-exclusion chromatography using a Superose 6 Increase 10/300 GL column equilibrated with Buffer A. Peak fractions were concentrated to ∼7 mg/ml with an Amicon Ultra concentrator with a 100-kDa molecular-weight cutoff and used to prepare cryo-EM grids.

The purification of WT MscM and MscM ΔPB1 in KCl buffer was performed as described above but substituting NaCl for equivalent concentrations of KCl. Additionally, in the case of WT MscM, the cells were lysed by cryo-milling (5 x 3 minutes at 30 Hz) instead of using a microfluidizer.

### Cryo-EM specimen preparation and data collection

All grids were prepared using a Vitrobot Mark IV (Thermo Fisher Scientific) set at 4°C and 100% humidity. Aliquots of 3.5 μl of the samples were applied to glowed-discharged Quantifoil R1.2/1.3 Cu 400 mesh holey carbon grids (Electron Microscopy Sciences). After a 30-s delay, the grids were blotted for 2-3 s with a blot force of 0. Grids were plunged into liquid nitrogen-cooled ethane.

All samples were imaged in the Cryo-EM Resource Center at the Rockefeller University using a 300-kV Titan Krios G2 electron microscope (Thermos Fisher Scientific) operated with SerialEM and equipped with a K3 direct detector camera and a Gatan GIF filter with a slit width of 20 eV. Data were recorded in counting mode at a nominal magnification of 105,000x, corresponding to a calibrated pixel size of 0.847 Å at the specimen level, and using a defocus range of -0.8 to -2.0 µm.

For WT MscM in NaCl buffer, movies were collected at a dose rate of 27.6 e^−^/s/pixel. Exposures of 1.2 s were dose-fractionated into 40 frames of 0.03 s, resulting in 1.16 e^−^/Å^2^/frame and a total dose of 46.21 e^−^/Å^2^.

For MscM ΔPB1 in NaCl buffer, MscM G912S in NaCl buffer, WT MscM in KCl buffer, and MscM ΔPB1ΔΤΜ7e in NaCl buffer, movies were collected at a dose rate of 28 e^−^/s/pixel. Exposures of 1.2 s were dose-fractionated into 40 frames of 0.03 s, resulting in 1.17 e^−^/Å^2^/frame and a total dose of 46.84 e^−^/Å^2^.

Data collection parameters are summarized in Table S1.

### Cryo-EM data processing of WT MscM in NaCl buffer

A total of 41,447 movies were collected that were motion-corrected using Patch Motion Correction and initial CTF values were estimated using Patch CTF Estimation using cryoSPARC Live^38^. Curation of the motion-corrected micrographs based on CTF values yielded 36,386 micrographs. All subsequent image processing was performed in cryoSPARC, and in all 3D-processing steps 7-fold symmetry was imposed.

An initial subset of 1,000 micrographs was used to pick particles using blob picker, yielding 231,421 particles that were extracted into 450×450-pixel boxes, binned 5-fold, and subjected to 2D classification into 50 classes. 6 classes with averages showing the clearest structural features were combined, yielding 32,983 particles that were subjected to 2D classification into 50 classes. 11 classes with averages showing the clearest structural features were combined, yielding 10,211 particles that were used to train an initial Topaz model^39^. This Topaz model was used to pick particles from the same subset of micrographs, resulting in 108,636 particles that were extracted into 450×450-pixel boxes, binned 5-fold, and subjected to 2D classification into 50 classes. 13 classes with averages showing the clearest structural features were combined, yielding 35,405 particles that were subjected to 2D classification into 50 classes. 13 classes with averages showing the clearest structural features were combined, yielding 12,170 particles that were used to train a second Topaz model. This Topaz model was used to pick particles from all the micrographs, yielding 7,016,041 particles that were extracted into 450×450-pixel boxes, binned 5-fold, and subjected to 2D classification into 100 classes. 15 classes with averages showing the clearest structural features were combined, yielding 1,504,523 particles that were subjected to 2D classification into 100 classes. 6 classes with averages showing the clearest structural features were combined and used to generate 3 *ab-initio* models, of which one clearly showed MscM and was used. The same particles were used to generate 4 decoy maps by prematurely terminating the reconstruction job. Of the 100 2D classes, 24 classes with poor averages were discarded and the remaining 76 classes were combined, yielding 1,064,397 particles. These particles were subjected to two iterative rounds of heterogeneous refinement to remove poor particles. Each heterogeneous refinement was seeded with 4 copies of the MscM *ab-initio* model and the 4 decoy maps. The particles that were assigned to the MscM classes were combined and the process was repeated, yielding 361,722 particles from 2 classes with 3D maps showing clear features of MscM. The particles were extracted into 450×450-pixel boxes without binning and subjected to heterogeneous refinement into 6 classes. 3 of the resulting maps resolved features in the cytoplasmic domain, and the particles in these classes were combined, yielding 193,758 particles for further processing. One of these 3 classes showed density for the periplasmic domain, and the 77,059 particles in this class were also further processed by themselves.

The 193,758 particles from the three classes with averages showing clear features for the cytoplasmic domain were subjected to non-uniform refinement, yielding a map at a resolution of 3.5 Å. To improve density for the cytoplasmic domain, the particles were subjected to two iterative rounds of 3D classification without alignment, first into 6 and then 4 classes, using a mask that included only the cytoplasmic domain and TMs 9-11. This process yielded a class with 50,627 particles with well-resolved density. The particles were subjected to local refinement, yielding a map at a resolution of 3.5 Å. After local and global CTF refinement, followed by another round of local refinement, the final map had a resolution of 3.4 Å according to gold-standard Fourier shell correlation (FSC) with a cut-off criterion of FSC = 0.143.

The 77,059 particles from the class showing density for the periplasmic domain were subjected to non-uniform refinement, yielding a map at a resolution of 3.6 Å. To improve the density for the periplasmic and TM domains, the particles were subjected to 3D classification without alignment into 6 classes. The class with the strongest density for the periplasmic domain was selected, and the 10,378 particles in this class were subjected to non-uniform refinement, resulting in a map at 6.7-Å resolution that partially resolved helices in the TM domain but not in the periplasmic domain.

### Cryo-EM data processing of MscM ΔPB1 in NaCl buffer

A total of 77,369 movies were collected from three grids. The movies were motion-corrected using Patch Motion Correction and initial CTF values were estimated using Patch CTF Estimation using cryoSPARC Live. Curation of the motion-corrected micrographs based on CTF values yielded 69,789 micrographs. All subsequent image processing was performed in cryoSPARC, and in all 3D-processing steps 7-fold symmetry was imposed.

An initial subset of 227 micrographs was used to pick particles using blob picker, yielding 46,336 particles that were extracted into 450×450-pixel boxes, binned 5-fold, and subjected to 2D classification into 50 classes. 11 classes with averages showing the clearest structural features were combined, yielding 10,839 particles that were used to train an initial Topaz model. This Topaz model was used to pick particles from ∼600 micrographs, yielding 75,939 particles that were extracted, binned, and subjected to 2D classification into 50 classes. 13 classes with averages showing the clearest structural features were combined, yielding 26,055 particles that were subjected to another round of 2D classification into 50 classes. 11 classes with averages showing the clearest structural features were combined, yielding 10,937 particles that were used to train a second Topaz model. This Topaz model was used for particle picking from ∼3,400 micrographs, yielding 414,131 particles that were extracted, binned, and subjected to 2D classification into 50 classes. 8 classes with averages showing the clearest structural features were combined, yielding 95,119 particles that were subjected to 2D classification into 50 classes. 14 classes with averages showing the clearest structural features were combined, yielding 43,331 particles that were used to train a third Topaz model.

This third Topaz model was used to pick particles from all the micrographs, yielding 11,459,225 particles that were extracted into 450×450-pixel boxes, binned 5-fold, and subjected to 2D classification into 100 classes. 7 classes with averages showing the clearest structural features were combined, yielding 1,589,457 particles that were subjected to another round of 2D classification into 100 classes. 8 classes with the averages showing the clearest structural features were combined and used to generate 3 *ab-initio* models, of which one clearly showed MscM and was used. The same particles were used to generate 4 decoy maps by prematurely terminating the reconstruction job. Of the 100 2D classes, 10 classes with poor averages were discarded and the remaining 90 classes were combined, yielding 10,774,331 particles. These particles were subjected to 6 iterative rounds of heterogeneous refinement to remove poor particles. Each heterogeneous refinement was seeded with 4 copies of the MscM *ab-initio* model and the 4 decoy maps. The particles that were assigned to the MscM classes were used to seed the subsequent round of heterogeneous refinement. This process yielded 1,590,870 particles, which were subjected to heterogeneous refinement seeded with 6 copies of the MscM *ab-initio* model to identify particles showing features for the periplasmic and the entire TM domain. Only one class showed features for all MscM domains. The 392,459 particles of this class were extracted into 450×450-pixel boxes without binning and subjected to non-uniform refinement, which yielded a map at 4.1-Å resolution, followed by local refinement, which improved the resolution of the map to 4.0 Å. These particles were subjected to two separate 3D classifications into 6 classes without alignment.

The first 3D classification used a mask that excluded the cytoplasmic domain and yielded one class of 58,387 particles that showed density for all periplasmic and TM helices. These particles were subjected to non-uniform refinement, and local and global refinement of the CTF values of the particles. A final non-uniform refinement yielded a map at a resolution of 4.3 Å, according to gold-standard FSC.

The second 3D classification used a mask that included only TMs 9-11 and the cytoplasmic domain and yielded one class of 135,601 particles with well-resolved features. After local refinement, the particles were classified again into 4 classes, resulting in one class of 61,183 particles showing the best resolved features. These particles were subjected to local refinement, and local and global refinement of the CTF values of the particles. A final local refinement yielded a map at a resolution of 3.4 Å, according to gold-standard FSC.

### Cryo-EM data processing of MscM G912S in NaCl buffer

A total of 10,906 movies were collected that were motion-corrected using MotionCorr2^40^, and initial CTF values were estimated with Patch CTF Estimation using cryoSPARC. Curation of the motion-corrected micrographs based on CTF values yielded 10,391 micrographs. All subsequent image processing was performed in cryoSPARC, and in all 3D-processing steps 7-fold symmetry was imposed.

An initial subset of 688 micrographs was used to pick particles using blob picker, yielding 104,642 particles that were extracted into 450×450-pixel boxes, binned 5-fold, and subjected to 2D classification into 50 classes. 4 classes with averages showing the clearest structural features were combined, yielding 12,249 particles that were submitted to another round of 2D classification into 50 classes. 11 classes with averages showing the clearest structural features were combined, yielding 5,444 particles that were used to train a Topaz model. This Topaz model was used to pick particles from all the micrographs, yielding 1,589,420 particles that were extracted into 450×450-pixel boxes, binned 5-fold, and subjected to 2D classification into 100 classes. 9 classes with averages showing the clearest structural features were combined, yielding 245,982 particles that were subjected to another round of 2D classification into 100 classes. 10 classes with averages showing the clearest structural features were combined, yielding 46,526 particles, that were used to generate 3 *ab-initio* models, of which one clearly showed MscM and was used. 30 classes from the same 2D classification with averages showing the clearest structural features were combined, yielding 182,037 particles that were subjected to heterogeneous refinement seeded with six copies of the *ab-initio* model.

### Cryo-EM data processing of WT MscM in KCl buffer

Initially, 10,309 movies were collected, motion-corrected using Patch Motion Correction and initial CTF values were estimated using Patch CTF Estimation using cryoSPARC Live. Curation of the motion-corrected micrographs based on CTF values yielded 8,745 micrographs. All subsequent image processing was performed in cryoSPARC, and in all 3D-processing steps 7-fold symmetry was imposed unless noted otherwise.

157 particles representing top and side views of MscM were interactively picked from 28 micrographs and used to train an initial Topaz model. This Topaz model was used to pick particles from 2,677 micrographs, yielding 101,758 particles that were extracted into 450×450-pixel boxes, binned 5-fold, and subjected to 2D classification into 100 classes. 16 classes with averages showing the clearest structural features were combined, yielding 29,868 particles.

These particles were used to train a second Topaz model. This Topaz model was used to pick particles from the same 2,677 micrographs, yielding 289,210 particles. Following two rounds of 2D classification into 100 classes, 9 classes with averages showing the clearest structural features were combined and used to generate 3 *ab-initio* models, of which one clearly showed MscM and was used. The same particles were used to generate 4 decoy maps by prematurely terminating the reconstruction job. The Topaz model was then used to pick particles from all the micrographs, yielding 1,085,918 particles that were extracted into 450×450-pixel boxes and binned 5-fold.

Two iterative rounds of 2D classification into 100 classes were performed, rejecting classes with averages showing poor features, yielding 312,607 particles. These particles were re-extracted into 450×450-pixel boxes without binning and subjected to heterogeneous refinement that was seeded with 4 copies of the MscM *ab-initio* model and the 4 decoy maps. The refinement yielded maps of MscM in two different conformations that were characterized by a flat or a curved TM domain. These classes were combined, and the resulting 256,043 particles were subjected to heterogeneous refinement that was seeded with three copies each of MscM with a curved and a flat TM domain. Two classes with the best resolved features for MscM with a curved TM domain were combined (97,364 particles) and one class showed the best resolved features for MscM with a flat TM domain (63,301 particles). These two particle stacks were separately subjected to non-uniform refinement, yielding maps at resolutions of 3.6 Å and 3.3 Å, respectively.

An additional 45,061 movies were collected, motion-corrected using Patch Motion Correction and initial CTF values were estimated using Patch CTF Estimation using cryoSPARC Live. Curation of the motion-corrected micrographs based on CTF values yielded an additional 38,262 micrographs that were added to the previous micrographs, yielding a total of 47,007 micrographs. The final Topaz model was used to pick particles from all the micrographs, yielding 5,857,401 particles that were extracted into 450×450-pixel boxes, binned 5-fold and subjected to three iterative rounds of 2D classification, rejecting classes with averages showing poor features. The remaining 1,968,140 particles were subjected to heterogeneous refinement that was seeded with two copies each of the MscM map with a curved TM domain and the MscM map with a flat TM domain that were obtained by non-uniform refinement and the 4 decoy maps. The 1,425,134 particles that were assigned to the MscM maps were re-extracted into 450×450-pixel boxes without binning and subjected to heterogeneous refinement with two copies each of the MscM maps with a flat and a curved TM domain. One class showing MscM with a flat TM domain showed high-resolution features, and the 450,036 particles in this class were subjected to non-uniform refinement, yielding a map at 3.1-Å resolution.

To improve the density for the TM domain, the particles were subjected to a 3D classification without alignment into 10 classes, using a mask including only the TM domain of MscM. Three classes yielded maps with density for all TM helices. The 136,448 particles from these classes were subjected to local refinement, and local and global refinement of the CTF values of the particles. A final local refinement yielded a map at a resolution of 2.8 Å, according to gold-standard FSC (map 1).

The map did not clearly resolve TMs 1-6. Therefore, the particles were symmetry expanded to C7 symmetry and subjected to a 3D classification without alignment using a mask including only the TM domain of a single subunit. One class showed a map with density for all TM helices. The 135,809 particles from this class were subjected to local refinement, and local and global refinement of the CTF values of the particles. A final local refinement yielded a map at a resolution of 3.2 Å, according to gold-standard FSC (map 2).

The initial map obtained with 450,036 particles did not show convincing density for the periplasmic domain. Therefore, the particles were subjected to three iterative rounds of heterogeneous refinement that were seeded with 6 copies of the map, rejecting after each round classes that yielded maps without density for the periplasmic domain. This process yielded the final 69,165 particles that were subjected to non-uniform refinement, yielding a map at a resolution of 3.3 Å (map 3) that revealed discernable, albeit low-resolution, density for the proximal periplasmic helical bundle PB2.

A composite map was created using maps 1-3 described above using UCSF ChimeraX^41^. First, map 2 was duplicated seven times and each copy was fit into the density representing a different subunit of map 1. An initial composite map (map 4) was created with map 1 and the seven copies of map 2 using the ‘volume maximum’ command. Map 3 was also fit into map 1 and density segmentation was performed on map 3 to isolate the density corresponding to the periplasmic domain. The density for the periplasmic domain was then joined with map 4 using the ‘volume maximum’ command to create a final composite map with density for all MscM domains.

### Cryo-EM data processing of MscM ΔPB1 in KCl buffer

A total of 11,623 movies were collected, motion-corrected using MotionCorr2, and initial CTF values were estimated with Patch CTF Estimation using cryoSPARC. Curation of the motion-corrected micrographs based on CTF values yielded 11,022 micrographs. All subsequent image processing was performed in cryoSPARC, and in all 3D-processing steps 7-fold symmetry was imposed.

127 particles representing top and side views of MscM were interactively picked from 18 micrographs and used to train an initial Topaz model. This Topaz model was used to pick particles from 1,459 micrographs, yielding 119,403 particles that were extracted into 450×450-pixel boxes, binned 5-fold, and subjected to 2D classification into 50 classes. 13 classes with averages showing the clearest structural features were combined, yielding 40,999 particles.

These particles were subjected to another round of 2D classification into 50 classes. 37 classes with averages showing the clearest structural features were combined, yielding 34,364 particles. These particles were used to train a second Topaz model, to generate 3 *ab-initio* models, of which one clearly showed MscM and was used, and to generate 4 decoy maps by prematurely terminating the reconstruction job. The new Topaz model was used to pick particles from all the micrographs, yielding 1,298,131 particles that were extracted into 450×450-pixel boxes, binned 5-fold, and subjected to 2D classification into 100 classes. 32 classes with the clearest structural features were combined, yielding 553,711 particles that were subjected to another round of 2D classification into 100 classes. 59 classes with the clearest structural features were combined, yielding 346,727 particles. These particles were subjected to 3 iterative rounds of heterogeneous refinements to remove poor particles. Each heterogeneous refinement was seeded with 4 copies of the MscM *ab-initio* model and the 4 decoy maps. The particles that were assigned to the MscM classes were used to seed the subsequent round of heterogeneous refinement. This process yielded 169,300 particles and showed maps of MscM with curved and flat TM domains. The particles were subjected to heterogeneous refinement that was seeded with 3 copies of MscM ΔPB1 in the closed conformation and three copies of WT MscM in the open conformation. The refinement yielded one class with well-resolved structural features of MscM with a flat TM domain. These 34,087 particles were re-extracted into 450×450-pixel boxes without binning and subjected to non-uniform refinement with on-the-fly local and global CTF refinement, yielding a map at a resolution of 3.2 Å, according to gold-standard FSC.

### Cryo-EM data processing of MscM ΔPB1ΔTM7e in NaCl buffer

A total of 26,496 movies were collected, motion-corrected using MotionCorr2, and initial CTF values were estimated with Patch CTF Estimation using cryoSPARC. Curation of the motion-corrected micrographs based on CTF values yielded 23,952 micrographs. All subsequent image processing was performed in cryoSPARC, and in all 3D-processing steps 7-fold symmetry was imposed.

An initial subset of 657 micrographs was used to pick particles using blob picker, yielding 110,556 particles that were extracted into 450×450-pixel boxes, binned 5-fold, and subjected to 2D classification into 50 classes. 6 classes showing averages with the clearest structural features were combined, yielding 20,751 particles that were used to train a Topaz model. This Topaz model was used to pick particles from all the micrographs, yielding 4,428,810 particles that were extracted into 450×450-pixel boxes, binned 5-fold, and subjected to 2D classification into 100 classes. 24 classes with well-resolved structural features were combined, yielding 1,591,986 particles that were subjected to another round of 2D classification into 100 classes. 66 classes with the clearest structural features were combined, yielding 1,341,261 particles. These particles were submitted to heterogeneous refinement that was seeded with 8 copies of the final map of MscM ΔPB1, yielding one class resolving the periplasmic, TM and cytoplasmic domains of MscM. These 253,534 particles were re-extracted into 450×450-pixel boxes without binning and subjected to non-uniform refinement, which yielded a map at 4.5-Å resolution. Two separate image-processing pathways were then performed.

To improve the density for the periplasmic and TM domains, the particles were subjected to 3D classification without alignment into 6 classes using a mask that omitted the cytoplasmic domain. One of the maps showed density for all periplasmic and TM helices, and the corresponding 48,883 particles were subjected to non-uniform refinement, yielding a map at 5.1 Å, according to gold-standard FSC.

To improve density for the cytoplasmic domain, the particles were first subjected to local refinement using a mask including only the cytoplasmic domain and TMs 9-11, which improved the resolution of the map to 3.9 Å. The particles were then subjected to 3D classification without alignment into 6 classes using the same mask. One of the maps showed the best density for the cytoplasmic domain and TMs 9-11, and the corresponding 57,877 particles were subjected to local refinement, and local and global refinement of the CTF values of the particles. A final local refinement yielded a map at a resolution to 3.5 Å, according to gold-standard FSC.

### Model building and refinement

Prior to model building, all cryo-EM maps were sharpened with EMReady^42^. To build an atomic model into the density map of WT MscM in NaCl buffer, the AlphaFold model for one MscM subunit was docked as a rigid body into the final local refinement map in UCSF ChimeraX^41^.

Due to the poor density for the entire periplasmic domain, the lack of density for TMs 1-6, and the discontinuous density for TMs 7-10, none of these regions were modeled. An atomic model was only built for TM11 and the cytoplasmic domain. This model was refined using real-space refinement in Coot^43^. The model for one monomer was further refined in Phenix^44^ (phenix.real_space_refine). 7 copies of the refined monomer were docked into the cryo-EM map using Phenix (phenix.dock_in_map), followed by a final round of refinement in Phenix (phenix.real_space_refine) to resolve any clashes between monomers.

To build an atomic model into the composite map of WT MscM in the open conformation, the AlphaFold model for one MscM subunit was docked as a rigid body into the map in UCSF ChimeraX. The monomer was refined using real-space refinement in Coot. The map showed no density for PB1 and the TM7-8 loop, which were therefore deleted from the model. Because the density for PB2 had low resolution, the AlphaFold model of PB2 was docked into the map as a rigid body. PB2 and TMs 1-6 were only modeled as poly-alanine peptide chains due to the poor resolution of those regions. Side chains were modeled for TMs 7-11 and the cytoplasmic domain. The protein model for one monomer was further refined in Phenix (phenix.real_space_refine) against the composite map. 7 copies of the refined monomer were docked into the composite map using Phenix (phenix.dock_in_map), followed by a final round of refinement in Phenix (phenix.real_space_refine) to resolve any clashes between monomers.

To build an atomic model into the density map of MscM ΔPB1 in NaCl buffer, the AlphaFold model for one MscM subunit was docked as a rigid body into the global refinement map of MscM ΔPB1 in UCSF ChimeraX. PB1 was deleted from the model. The monomer was refined using real-space refinement in Coot using the global refinement map for PB2 and TMs 1-10 and the local refinement map for TM11 and the cytoplasmic domain. PB2 and TMs 1-10 were only modeled as poly-alanine peptide chains due to the poor resolution of those regions. Density for the TM7-8 loop was missing, and the loop was thus deleted from the model. Side chains were modeled for TM11 and the cytoplasmic domain. The protein model for one monomer was further refined in Phenix (phenix.real_space_refine) using the global refinement map. 7 copies of the refined monomer were docked into the global refinement map using Phenix (phenix.dock_in_map), followed by a final round of refinement in Phenix (phenix.real_space_refine) using the local refinement map to resolve any clashes between monomers in TM11 and the cytoplasmic domain only. The same procedure was used to create a model for MscM ΔPB1ΔTM7e, but here an atomic model could be built for TMs 8 and 9.

All refinement and validation statistics were determined using Phenix (phenix.validation_cryoem) and are listed in Table S1. For the structure of MscM ΔPB1 in NaCl buffer, statistics were calculated for both the entire model using the global refinement map, and TM11 and the cytoplasmic domain using the local refinement map. For the structure of MscM in the open conformation, statistics were calculated excluding PB2 due to its poor resolution.

Statistics were calculated for both a single subunit using the symmetry-expanded locally refined map, and the entire model using the composite map. For the structure of MscM ΔPB1ΔTM7e in NaCl buffer, statistics were calculated for both the entire model using the global refinement map, and TMs 9-11 and the cytoplasmic domain using the local refinement map.

### Giant spheroplasts preparation

The *E. coli* strain MJF431 was used for preparation of giant spheroplasts. 4 ml LB medium supplemented with 100 μg/ml ampicillin was inoculated with colonies of *E. coli* carrying pTRC99a vector with WT or mutant MscM. The overnight cultures were used to seed fresh LB medium (30 ml) supplemented with 100 μg/ml ampicillin. The cells were grown for ∼1.5 hours at 37°C and when OD_600_ reached ∼0.5, 3 ml of the culture was used to inoculate 27 ml of fresh LB medium supplemented with 100 μg/ml ampicillin and 60 μg/ml cephalexin. The cells were grown for 1.5 hours at 37°C to allow the formation of long, filamentous cells. Protein expression was induced with 1 mM IPTG and allowed to proceed for another 1.5 hours at 37°C. The cells were harvested by centrifugation at 3,000xg for 5 minutes at room temperature, washed with 2.5 ml of 0.8 M sucrose in deionized water, pelleted again and resuspended in 2.5 ml of fresh 0.8 M sucrose in deionized water. Spheroplast formation was initiated by the addition of 150 μl of 1 M Tris-HCl, pH 8.0, 120 μl 5 mg/mg lysozyme, 50 μl of 5 mg/ml DNase I and 150 μl of 125 mM EDTA, pH 8.0. After 15 minutes of incubation at room temperate, the reaction was quenched by the addition of 1 ml of stop solution (10 mM Tris-HCl, pH 8.0, 20 mM MgCl_2_, 0.8 M sucrose). The spheroplasts were precipitated at 800xg for ∼5-8 minutes at 4°C over a cushion solution (10 mM Tris-HCl, pH 8.0, 10 mM MgCl_2_, 0.8 M sucrose). The spheroplasts were resuspended in cushion solution and stored at -20°C until further use.

### Patch-clamp electrophysiology

The bath and pipette solution used for recordings contained 5 mM HEPES, pH 7.2, 200 mM KCl, 90 mM MgCl_2_ and 2 mM CaCl_2_, with the bath solution also supplemented with 400 mM sucrose to stabilize the integrity of the spheroplasts. The currents were amplified with an Axopatch 200B amplifier (Molecular Devices), filtered at 1 kHz, and data were acquired at either 5 or 20 kHz with a Digidata 1550B digitizer using pCLAMP 11.4.3 acquisition software (Molecular Devices). Negative pressure was applied to the recording pipette using a High-Speed Pressure Clamp-1 (ALA Scientific Instruments). The membrane potential was held constant at +30 mV for all recordings.

The activation-pressure ratio for MscM is defined as the ratio between the pressure at which the first MscL channel opens and the pressure at which the first MscM channel opens. The hysteresis ratio for MscM is defined as the ratio of the mid-point opening pressure to the mid-point closing pressure of MscM. All recordings that were used to calculate the activation-pressure ratio and hysteresis of MscM were performed in three technical replicates. For each patch, the reported values represent the average from the technical replicates.

The time constant of activation used to characterize the activation kinetics of MscM was determined by fitting exponential curves to the pressure pulse recordings in ClampFit.

Open-dwell times were calculated from several recordings from different patches for each construct using a single-channel detection threshold approach.

Recordings were visualized with ClampFit, and all graphs were made with GraphPad Prism. To reduce the noise in the electrophysiological recordings for visualization, a low-pass Gaussian filter was applied with 27 coefficients.

### MD simulations

Systems for all-atom MD simulations of MscS and the core structure of MscM (a.a. 830 – 1085) in the closed and open conformations, MscM ΔPB1 and MscM ΔPB1 ΔPBTM7 respectively, were built using the CHARMM-GUI Membrane Builder web server^45,46^. Each channel was placed in a membrane composed of 1-palmitoyl-2-oleoyl-sn-glycero-3-phosphoethanolamine.

The TIP3P water model and 1 M KCl was used to solvate the system^47,48^. The van der Waals interactions were smoothly switched off between 10 and 12 Å using the force-switch method^49^. Long-range electrostatic interactions were calculated using the Particle Mesh Ewald method, with a cut-off value of 1.2 nm, a Fourier grid spacing of 0.12 nm, and an interpolation order of 4 for the Ewald mesh^50,51^. The v-rescale thermostat and C-rescale barostat were used to maintain the temperate and pressure at 310.15 K and 1 atm, respectively^52,53^. Simulations were run with a 2-fs time step. The LINCS algorithm was used to constrain hydrogen bonds^54^. The GROMACS 2025 software package and the CHARMM36m force field were used to run all simulations^55–57^.

The systems were energy minimized and equilibrated using the standard CHARMM-GUI protocols, with a 6-step equilibration protocol that gradually relaxes position restraints. The final equilibration step that restrains only the protein backbone was extended to a total of 100 ns to allow lipids to associate with the hydrophobic pockets of the channels. After this extended equilibration, restraint-free production runs were performed for another 1 μs. Three independent replica simulations were performed for each channel.

The number of ions in the cytoplasmic domain of the channel was calculated using the GROMACS command ‘gmx select.’ An ion was considered to be in the cytoplasmic domain if it was within 2.8 nm of the center of mass (COM) of the cytoplasmic domain (MscS a.a. 127-276; MscM a.a. 930-1075). The permeability events were calculated using the output list of the unique residue identities of ions in the cytoplasmic domain generated with the ‘gmx select’ command and a custom bash script generated with ChatGTP.

Any ion that entered the cytoplasmic domain and remained within 2.8 nm of the COM for at least 3 ns was counted as an entry event. Any ion that exited the cytoplasmic domain and remained further away than 2.8 nm from the COM for at least 3 ns was considered as an exit event.

All graphs were made in GraphPad Prism. Simulation snapshots and movies were generated in UCSF ChimeraX.

### *E. coli* growth curves

*E. coli* strain MJF641 (lacking all MS channels) was transformed with either WT MscM or MscM deletion mutants in a pTRC99a vector and grown on LB plates supplemented with 100 μg/ml ampicillin. Individual colonies were picked, used to inoculate 4 ml of M9 minimal medium supplemented with 0.4% (w/v) D-glucose and 100 μg/ml ampicillin, and cultures were grown overnight at 37°C. The overnight cultures were used to seed fresh M9 minimal medium cultures at an OD_600_ of ∼0.02. The cultures were incubated for 4 hours at 37°C, at which point the OD_600_ reached ∼0.1. Protein expression was induced by the addition of 100 μM IPTG. Cell growth was monitored for an additional 8 hours. Every hour, aliquots of the cultures were taken and the OD_600_ was measured using a NanoDrop 2000 (Thermo Fisher Scientific).

### Data availability

The cryo-EM maps have been deposited in the Electron Microscopy Data Bank under accession codes EMD-76005 (WT MscM in NaCl, closed conformation, global refinement), EMD-76007 (WT MscM in NaCl, closed conformation, local refinement), EMD-76009 (MscM ΔPB1 in NaCl, closed conformation, global refinement), EMD-76010 (MscM ΔPB1 in NaCl, closed conformation, local refinement), EMD-76011 (WT MscM in KCl, open conformation, global refinement), EMD-76013 (WT MscM in KCl, open conformation, symmetry expanded refinement), EMD-76015 (WT MscM in KCl, open conformation, global refinement with density for PB2), EMD-76016 (WT MscM in KCl, open conformation, composite structure), EMD-76017 (MscM ΔPB1 in KCl, open conformation), EMD-76018 (MscM ΔPB1ΔTM7e in NaCl, global refinement) and EMD-76019 (MscM ΔPB1ΔTM7e in NaCl, local refinement). The atomic coordinates have been deposited in the Protein Data Bank under accession codes 11SM (WT MscM in NaCl, closed), 11SO (MscM ΔPB1 in NaCl, closed), 11SQ (WT MscM in KCl, open) and 11SR (MscM ΔPB1ΔTM7e in NaCl, closed).

## ACKNOWLEDGEMENTS

We thank M. Ebrahim, J. Sotiris, and H. Ng at the Evelyn Gruss Lipper Cryo-EM Resource Center of The Rockefeller University for assistance with cryo-EM data collection and members of the Walz group for helpful discussions. We thank M. Bush for cloning the MscM gene from the *E. coli* genome, and E. Brown and R. MacKinnon for providing *P. pastoris* strain SMD1163H. This work was supported by National Institutes of Health grant R01 GM144581 (T.W.).

## Author contributions

T.W. conceived the study. G.H. and T.W. designed the experiments. G.H. performed all experiments. G.H. and T.W. analyzed the results. G.H. and T.W. wrote the manuscript.

