## Supplementary Information for "MscM uses a novel gating mechanism for bacterial mechanosensitive channels"

### SUPPLEMENTARY FIGURES

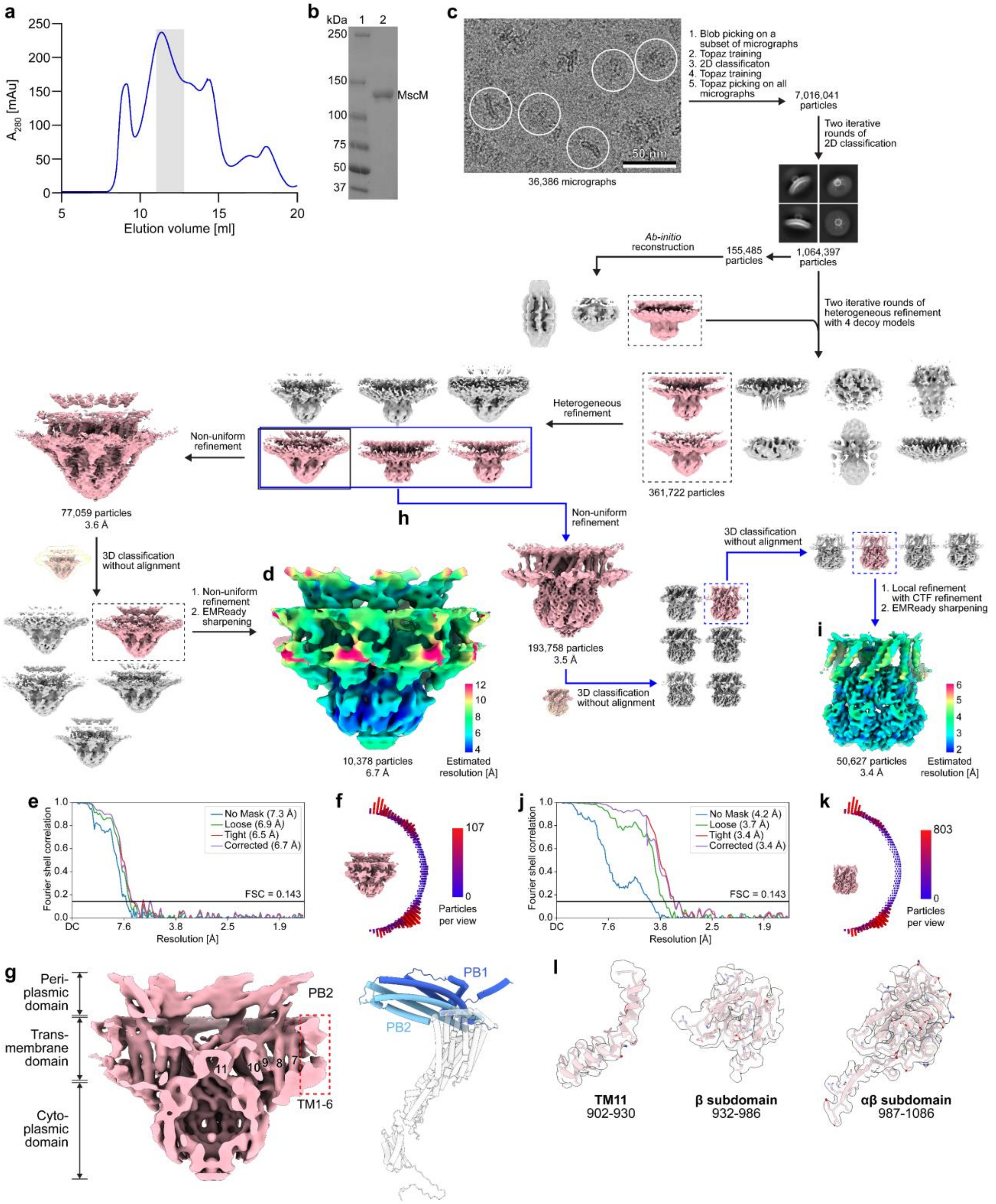

**Supplementary Figure 1. Purification and cryo-EM analysis of WT MscM in NaCl buffer. a** Size-exclusion chromatogram of MscM expressed in *P. pastoris* and purified in GDN. **b** Coomassie bluestained 12% SDS-PAGE gel. Lane 1: molecular weight markers; lane 2: pooled fractions indicated by the shaded region in panel a. **c** Raw image of vitrified WT MscM in NaCl buffer low-pass filtered to 2 Å with some representative particles circled and image-processing workflow used to generate a map of the entire protein (black arrows and dashed boxes). **d** Final map of WT MscM in NaCl buffer showing the channel in the closed conformation but missing density for peripheral TMs 1-6. The map is colored based on local-resolution estimates. **e** Gold-standard Fourier-shell correlation (FSC) curves. The resolution values indicated are based on the FSC = 0.143 cutoff criterion. **f** Angular distribution of the particles included in the final reconstruction. **g** Left panel: cut-away view of the final map with structural elements labeled. Right panel: AlphaFold model of a single MscM subunit shown with the periplasmic domain colored in blue, and the TM and cytoplasmic domains colored in white. **h** Image-processing workflow used to generate a map of the MscS-like core structure (blue arrows and dashed boxes). **i** Final map of the core structure of WT MscM in NaCl buffer colored based on local-resolution estimates. **j** Gold-standard FSC curves. The resolution values indicated are based on the FSC = 0.143 cutoff criterion. **k** Angular distribution of the particles included in the final reconstruction. **l** Cryo-EM density with the atomic model for TM11 and the β and αβ subdomains of the cytoplasmic domain. The numbers indicate the modeled amino-acid residues.

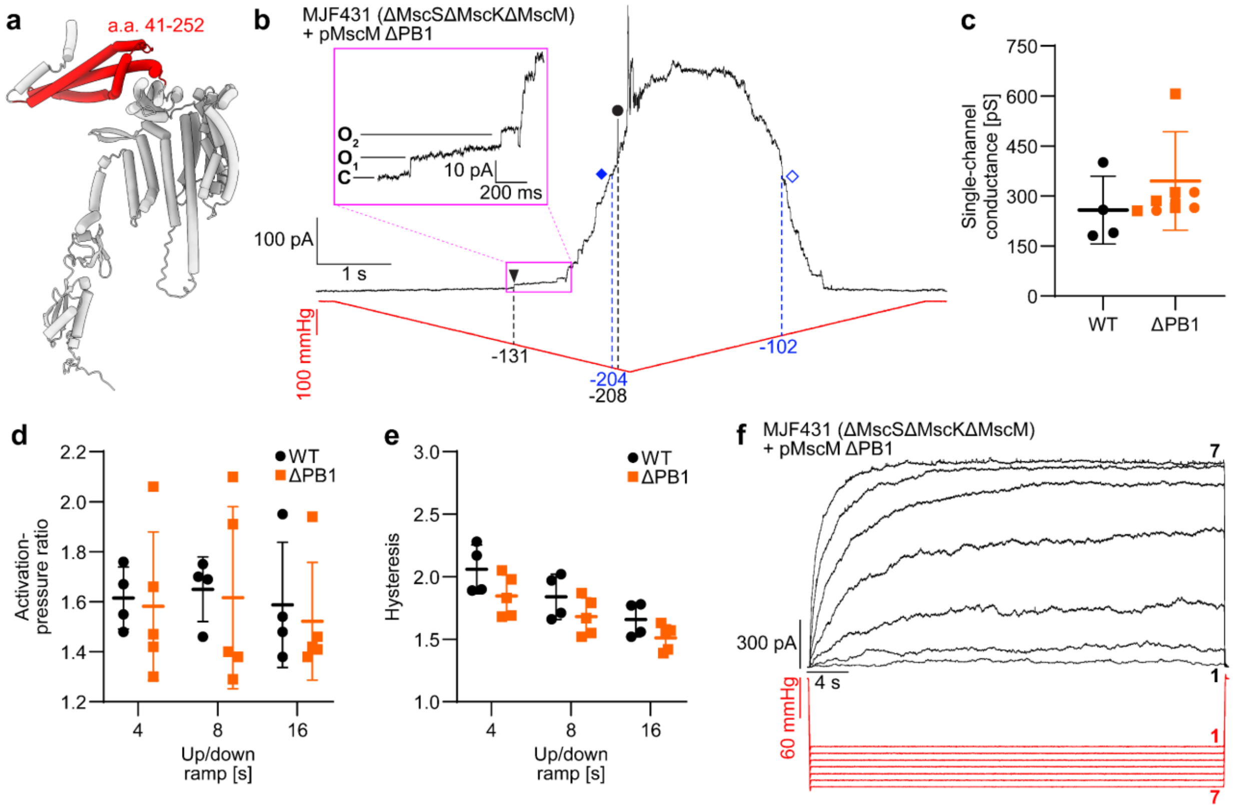

**Supplementary Figure 2. Electrophysiological characterization of the MscM ΔPB1 mutant. a** AlphaFold model of a single MscM subunit (white) indicating the membrane-distal helical bundle of the periplasmic domain, PB1 (red), that was deleted in the MscM ΔPB1 mutant. Note that the two helices preceding PB1 are formed by the signal sequence. **b** Representative patch-clamp recording (n = 5) from a giant spheroplast generated from *E. coli* strain MJF431 (which does not express MscS, MscK and MscM) overexpressing MscM ΔPB1 in response to a symmetric 8-s-long pressure ramp from 0 mmHg to -210 mmHg and back to 0 mmHg. Currents were recorded in the inside-out excised patch configuration with a pipette potential of +30 mV. Pressure applied to the patch membrane and the recorded current are shown in red and black, respectively. Black triangle: first opening of MscM; black circle: first opening of MscL; filled blue rhombus: mid-point activation of MscM; empty blue rhombus: mid-point deactivation of MscM; dashed black lines: pressures at which the first MscM and the first MscL channels open; dashed blue lines: pressures at which the mid-point activation and mid-point deactivation of MscM occur. The inset shows a zoomed-in view of the MscM currents in the region indicated by the purple box. C: current level when all MscM channels are closed; $O_1$-$O_2$: current levels when one or two MscM channels are open. **c** Quantitative analysis shows no significant change in the single-channel conductance of MscM ΔPB1 (orange squares) compared to that of WT MscM (black circles). **d** Quantitative analysis shows no significant change in the activation-pressure ratio of MscM ΔPB1 (orange squares) compared to that of WT MscM (black circles) in response to ramps of different speeds. **e** Quantitative analysis shows no significant change in the hysteresis ratio of MscM ΔPB1 (orange squares) compared to that of WT MscM (black circles) in response to ramps of different speeds. **f** Representative patch-clamp recording (n = 5) from a giant spheroplast generated from *E. coli* strain MJF431 overexpressing MscM ΔPB1 in response to 30-s-long pulses with pressure increasing from -100 mmHg to -160 mmHg in -10 mmHg intervals. The PB1 deletion does not alter the slow activation kinetics of MscM or its lack of desensitization/inactivation.

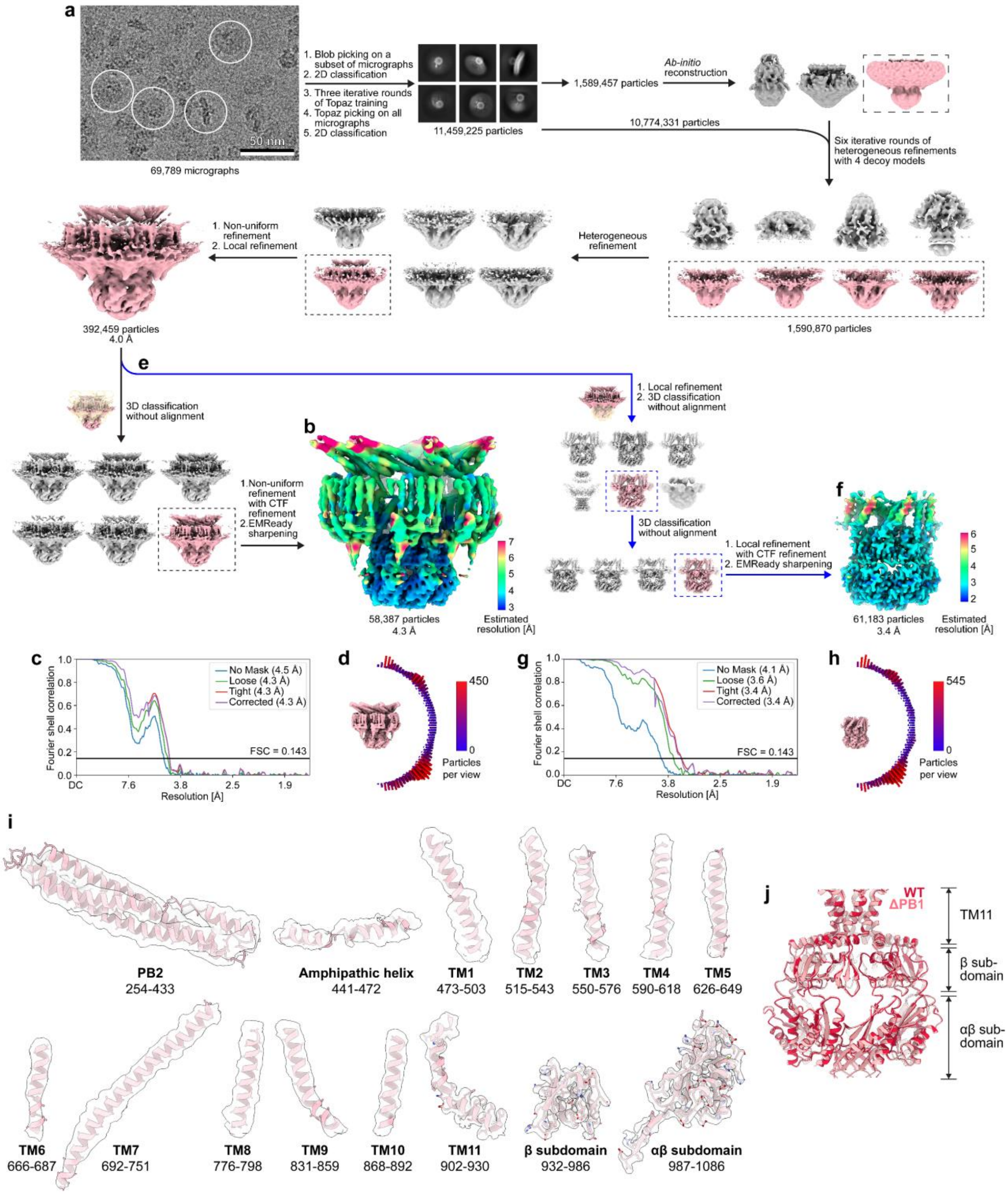

**Supplementary Figure 3. Cryo-EM analysis of MscM ΔPB1 in NaCl buffer**. **a** Raw image of vitrified MscM ΔPB1 in NaCl buffer low-pass filtered to 2 Å with some representative particles circled and image-processing workflow used to generate a map of the entire protein (black arrows and dashed boxes). **b** Final map of MscM ΔPB1 in NaCl buffer showing the channel in the closed conformation with density for all domains. The map is colored based on local-resolution estimates. **c** Gold-standard Fourier-shell correlation (FSC) curves. The resolution values indicated are based on the FSC = 0.143 cutoff criterion. **d** Angular distribution of the particles included in the final reconstruction. **e** Image-processing workflow used to generate a map of the MscS-like core structure (blue arrows and dashed boxes). **f** Final map of the core structure of WT MscM in NaCl buffer. **g** Gold-standard FSC curves. The resolution values indicated are based on the FSC = 0.143 cutoff criterion. **h** Angular distribution of the particles included in the final reconstruction. **i** Cryo-EM density with either backbone models for PB2, the amphipathic helix and TMs 1-10 or atomic models for TM11 and the β and αβ subdomains of the cytoplasmic domain. The numbers indicate the modeled amino-acid residues. **j** Alignment of TM11 and the cytoplasmic domain of WT MscM (crimson ribbons) and MscM ΔPB1 (pink ribbons) showing that the two structures are essentially identical.

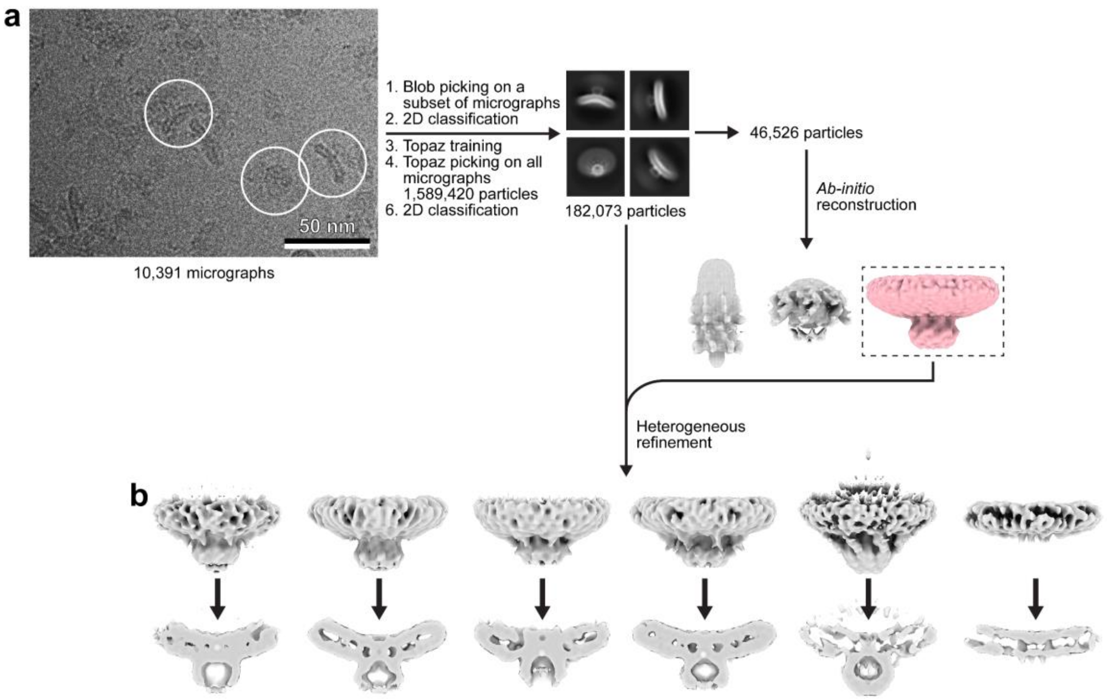

**Supplementary Figure 4. Cryo-EM analysis of MscM G912S in NaCl buffer**. **a** Raw image of vitrified MscM G912S in NaCl buffer with some representative particles circled and image-processing workflow. **b** Views parallel to the membrane (top panel) and cut-away views (bottom panel) of the maps resulting from the final heterogeneous refinement. All maps show curved TM domains, indicating that all the channels adopt a closed conformation.

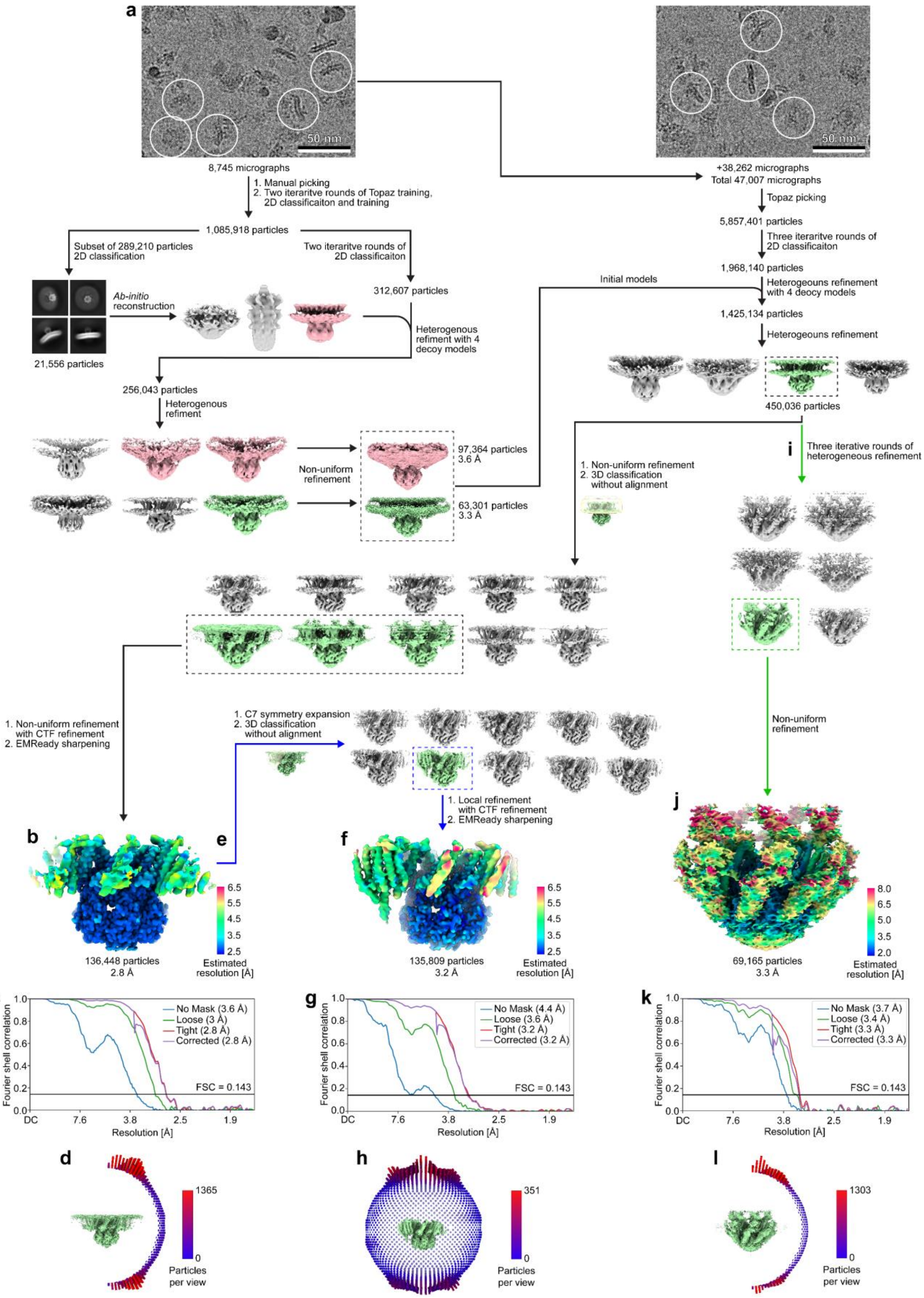
a
50 nm
50 nm
8,745 micrographs
1. Manual picking
2. Two iteraritve rounds of Topaz training, 2D classificaiton and training
1,085,918 particles
Subset of 289,210 particles 2D classification
Two iteraritve rounds of 2D classificaiton
312,607 particles
Ab-initio reconstruction
21,556 particles
Heterogenous refiment with 4 decoy models
256,043 particles
Heterogenous refiment
Non-uniform refinement
97,364 particles 3.6 Å
63,301 particles 3.3 Å
+38,262 micrographs
Total 47,007 micrographs
Topaz picking
5,857,401 particles
Three iteraritve rounds of 2D classificaiton
1,968,140 particles
Initial models
Heterogeouns refinement with 4 deocy models
1,425,134 particles
Heterogeouns refinement
450,036 particles
1. Non-uniform refinement
2. 3D classification without alignment
i
Three iterative rounds of heterogeneous refinement
1. Non-uniform refinement with CTF refinement
2. EMReady sharpening
1. C7 symmetry expansion
2. 3D classification without alignment
1. Local refinement with CTF refinement
2. EMReady sharpening
Non-uniform refinement
b
e
f
j
136,448 particles
2.8 Å
135,809 particles
3.2 Å
69,165 particles
3.3 Å
Estimated resolution [Å]
c
g
k
Fourier shell correlation
Resolution [Å]
No Mask (3.6 Å)
Loose (3 Å)
Tight (2.8 Å)
Corrected (2.8 Å)
No Mask (4.4 Å)
Loose (3.6 Å)
Tight (3.2 Å)
Corrected (3.2 Å)
No Mask (3.7 Å)
Loose (3.4 Å)
Tight (3.3 Å)
Corrected (3.3 Å)
FSC = 0.143
DC
7.6
3.8
2.5
1.9
d
h
l
1365
351
1303
Particles per view

**Supplementary Figure 5. Cryo-EM analysis of WT MscM in KCl buffer**. **a** Raw images of vitrified WT MscM in KCl buffer low-pass filtered to 2 Å with some representative particles circled and image-processing workflow used to generate a map of the entire protein (black arrows and dashed boxes). **b** Final map of WT MscM in KCl buffer showing the channel in the open conformation with low-resolution density for peripheral TMs 1-6 and missing density for the periplasmic domain. The map is colored based on local-resolution estimates. **c** Gold-standard Fourier-shell correlation (FSC) curves. The resolution values indicated are based on the FSC = 0.143 cutoff criterion. **d** Angular distribution of the particles included in the final reconstruction. **e** Image-processing workflow used to generate a map resolving all TM helices in one MscM subunit. **f** Map of WT MscM in KCl buffer resolving all the TM helices in one subunit and colored based on local-resolution estimates. **g** Gold-standard FSC curves. The resolution values indicated are based on the FSC = 0.143 cutoff criterion. **h** Angular distribution of the particles included in the reconstruction. **i** Image-processing workflow used to generate a map that shows density for the periplasmic domain. **j** Final map of WT MscM in KCl buffer showing density for the periplasmic domain and colored based on local-resolution estimates. **k** Gold-standard FSC curves. The resolution values indicated are based on the FSC = 0.143 cutoff criterion. **l** Angular distribution of the particles included in the final reconstruction.

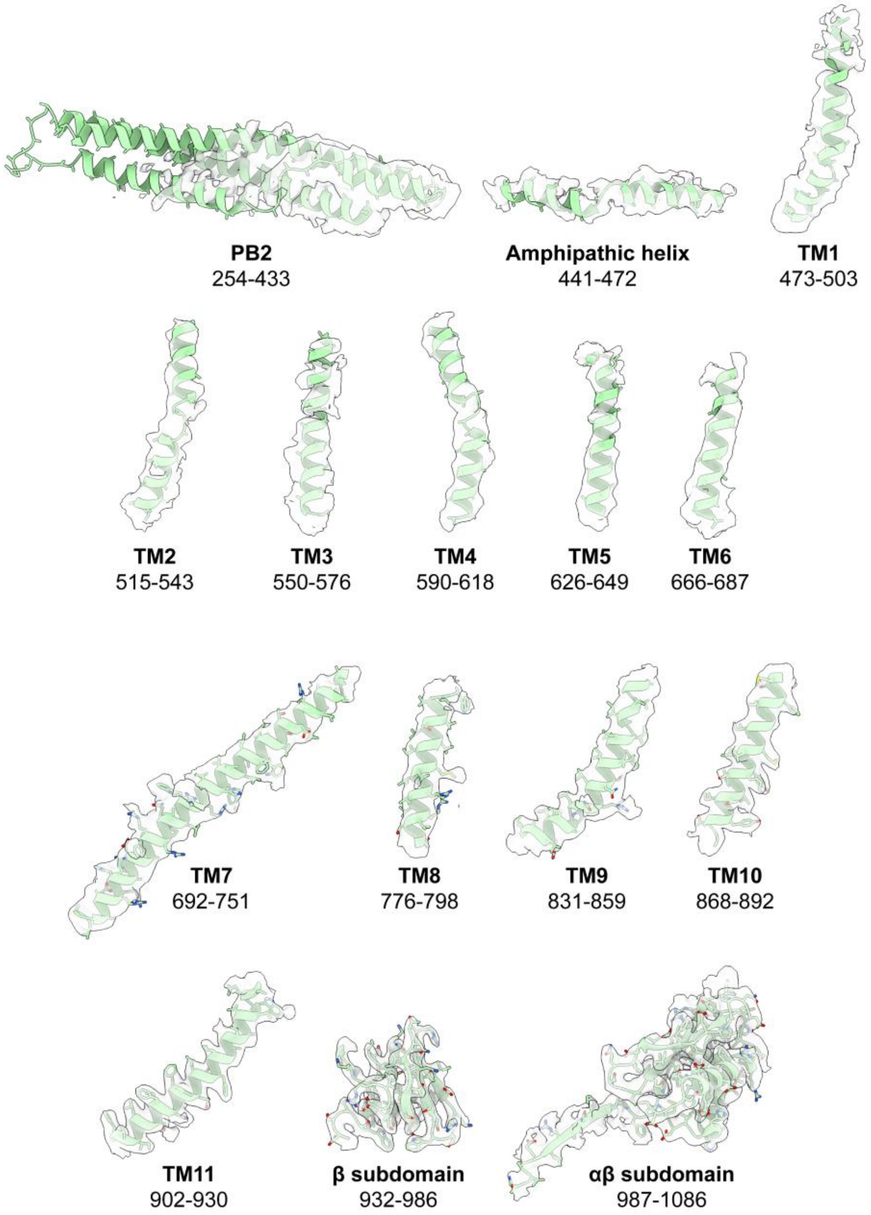

**Supplementary Figure 6. Map and model of WT MscM in KCl buffer.** Cryo-EM density with either backbone models for PB2, the amphipathic helix and TMs 1-6 or atomic models for TMs 7-11 and the β and αβ subdomains of the cytoplasmic domain. The numbers indicate the modeled amino-acid residues.

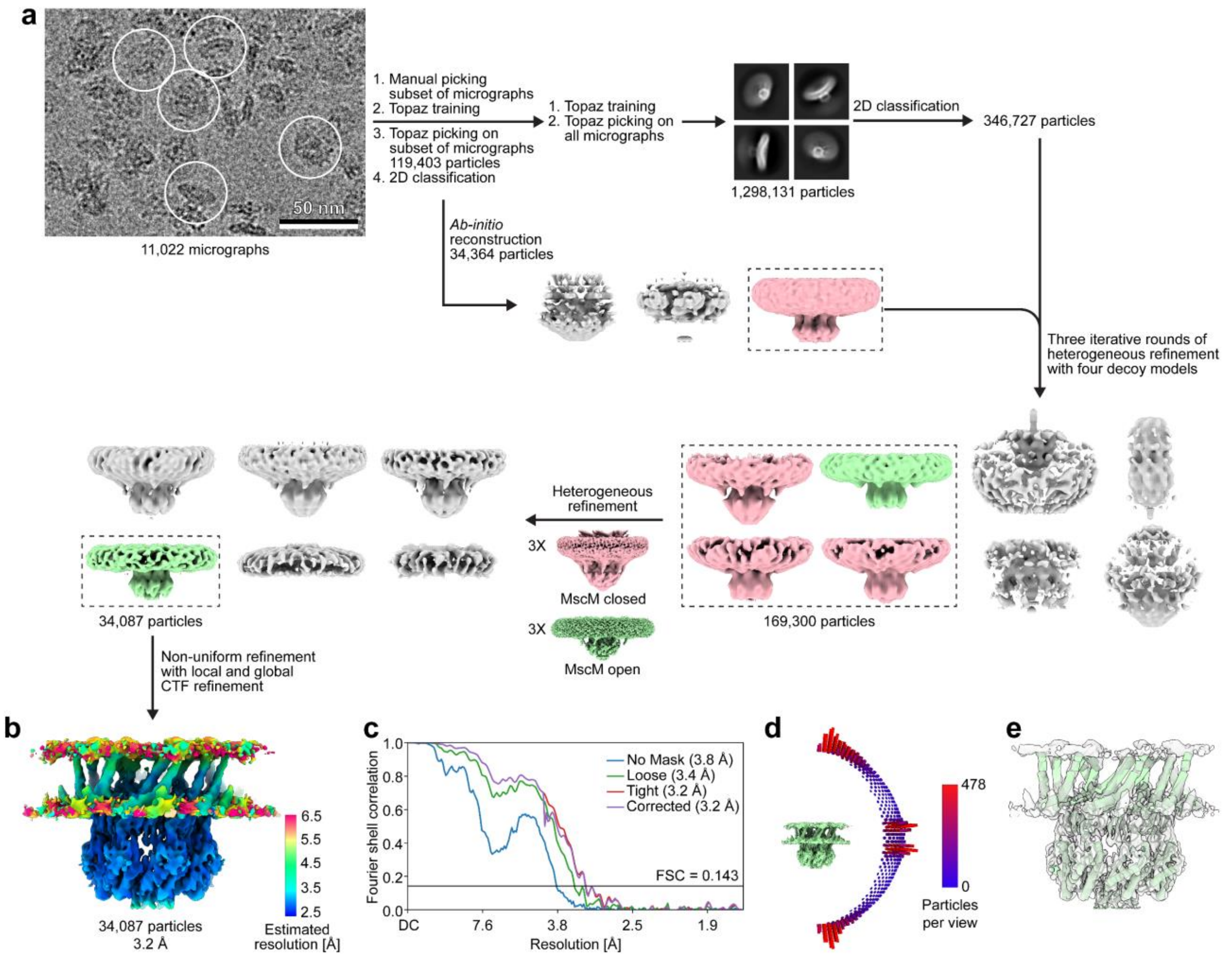

**Supplementary Figure 7. Cryo-EM analysis of MscM ΔPB1 in KCl buffer**. **a** Raw image of vitrified MscM ΔPB1 in KCl buffer low-pass filtered to 2 Å with some representative particles circled and image-processing workflow used to generate a map of the protein. **b** Final map of MscM ΔPB1 in KCl buffer showing the channel in the open conformation but missing density for the peripheral TMs 1-6. The map is colored based on local-resolution estimates. **c** Gold-standard Fourier-shell correlation (FSC) curves. The resolution values indicated are based on the FSC = 0.143 cutoff criterion. **d** Angular distribution of the particles included in the final reconstruction. **e** Fit of the model of MscM in the open conformation obtained from the cryo-EM analysis of WT MscM in KCl buffer (green cylinders) into the density map shown in panel b (white transparent surface).

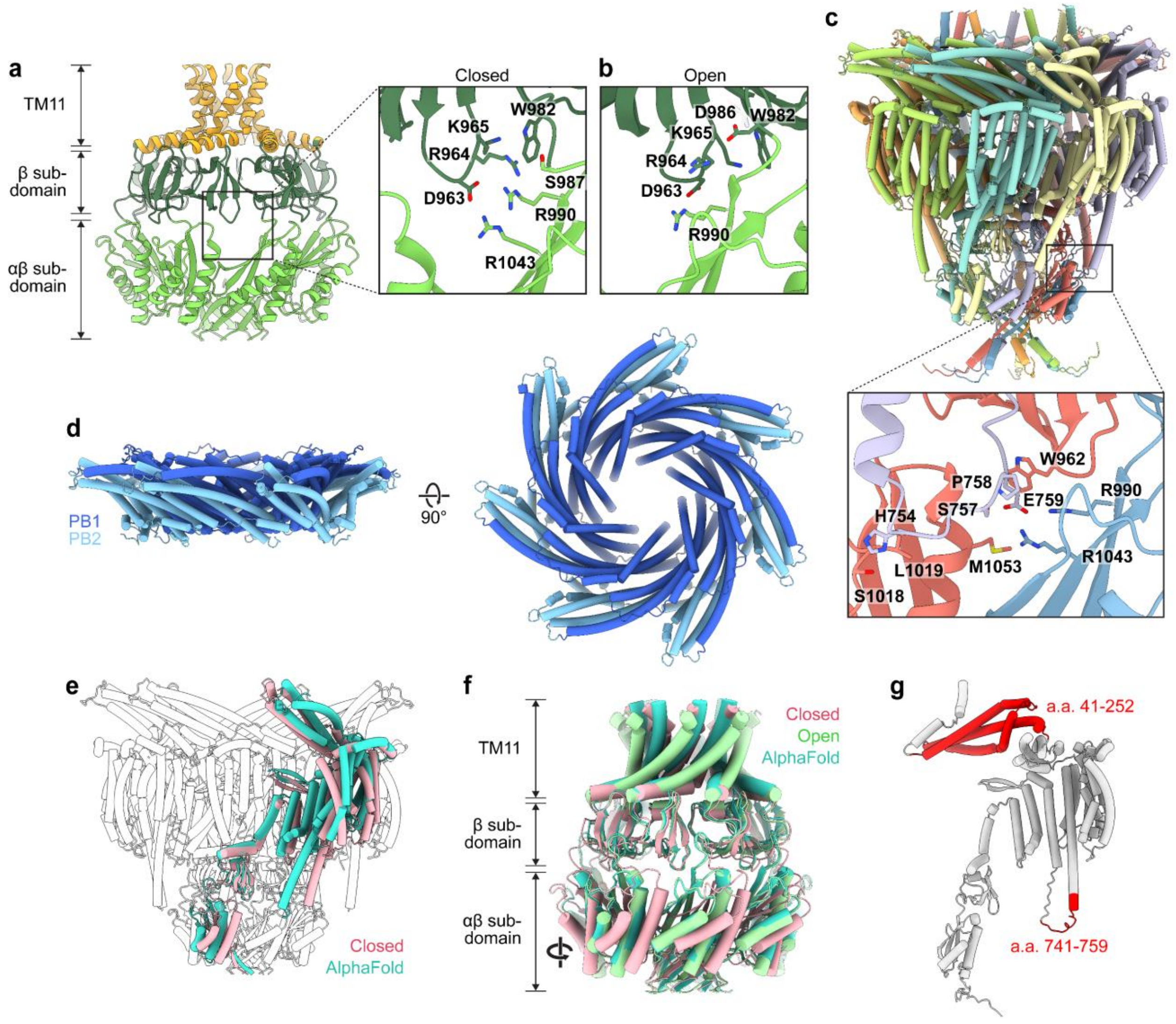

**Supplementary Figure 8. Structural analysis of the two conformations adopted by the cytoplasmic domain and AlphaFold prediction of the entire channel. a** Structure of TM11 (orange) and the cytoplasmic domain (β subdomain in dark green and αβ subdomain in light green) of MscM in the closed conformation. The inset shows a zoomed-in view of a fenestration with the residues stabilizing the closed conformation of the cytoplasmic domain shown as sticks and labeled. **b** Zoomed-in view of a fenestration of MscM in the open conformation with the residues stabilizing the open conformation of the cytoplasmic domain shown as sticks and labeled. **c** AlphaFold-Multimer prediction of an MscM channel shown in cartoon representation and colored according to subunit. The inset shows a zoomed-in view of the interactions between the TM7-8 loop and the cytoplasmic domain with the residues mediating these interactions shown as sticks and labeled. **d** Views parallel (left) and perpendicular (right) to the membrane of the MscM periplasmic domain structure (PB1 in dark blue and PB2 in light blue) as predicted by AlphaFold-Multimer, suggesting that PB1 forms a ring structure that is nestled into the ring formed by PB2. **e** Alignment of the AlphaFold prediction of MscM (one subunit shown in cyan) aligned to the closed conformation of MscM (one subunit shown in pink and six subunits shown in white). For clarity, the predicted structure of PB1 is not shown. **f** Alignment of the structure of TM11 and the cytoplasmic domain predicted by AlphaFold-Multimer (cyan) and those in the closed (pink) and open (green) conformations determined by cryo-EM. AlphaFold did not predict the novel conformation of the MscM cytoplasmic domain in the closed conformation. **g** AlphaFold model of a single MscM subunit (white) indicating the regions that were deleted (red) in the MscM ΔPB1ΔTM7e mutant.

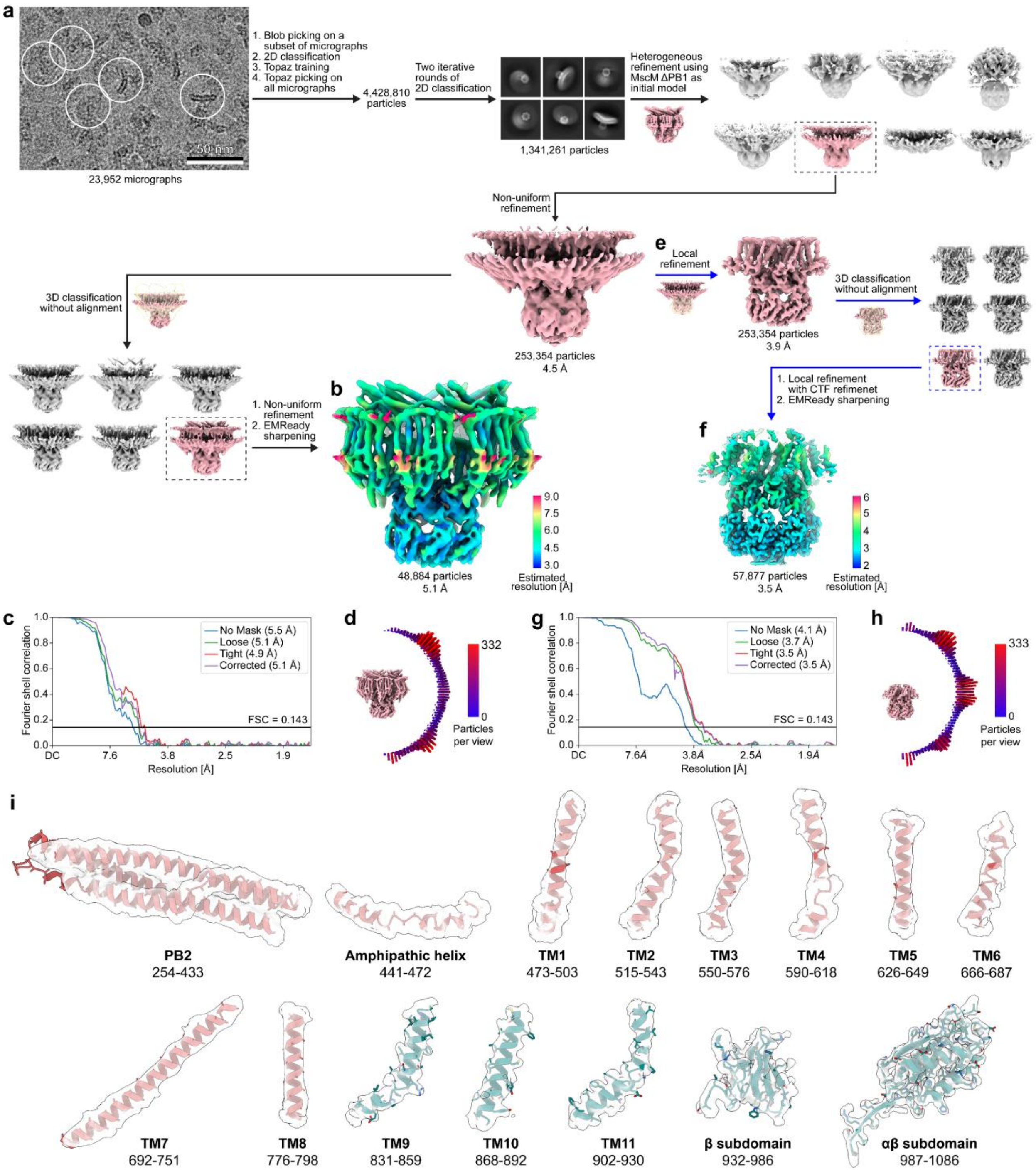

**Supplementary Figure 9. Cryo-EM analysis of MscM ΔPB1ΔTM7e in NaCl buffer**. **a** Raw image of vitrified MscM ΔPB1ΔTM7e in NaCl buffer low-pass filtered to 2 Å with some representative particles circled and image-processing workflow that was used to generate a map of the entire protein (black arrows and dashed boxes). **b** Final map of MscM ΔPB1ΔTM7e in NaCl buffer showing the channel in the closed conformation with density for all domains. The map is colored based on local-resolution estimates. **c** Gold-standard Fourier-shell correlation (FSC) curves. The resolution values indicated are based on the FSC = 0.143 cutoff criterion. **d** Angular distribution of the particles included in the final reconstruction. **e**

Image-processing workflow used to generate a map of the MscS-like core structure (blue arrows and dashed boxes). **f** Final map of the core structure of MscM ΔPB1ΔTM7e in NaCl buffer. **g** Gold-standard FSC curves. The resolution values indicated are based on the FSC = 0.143 cutoff criterion. **h** Angular distribution of the particles included in the final reconstruction. **i** Cryo-EM density with either backbone models for PB2, the amphipathic helix and TMs 1-8 or atomic models for TMs 9-11 and the β and αβ subdomains of the cytoplasmic domain. The numbers indicate the modeled amino-acid residues.

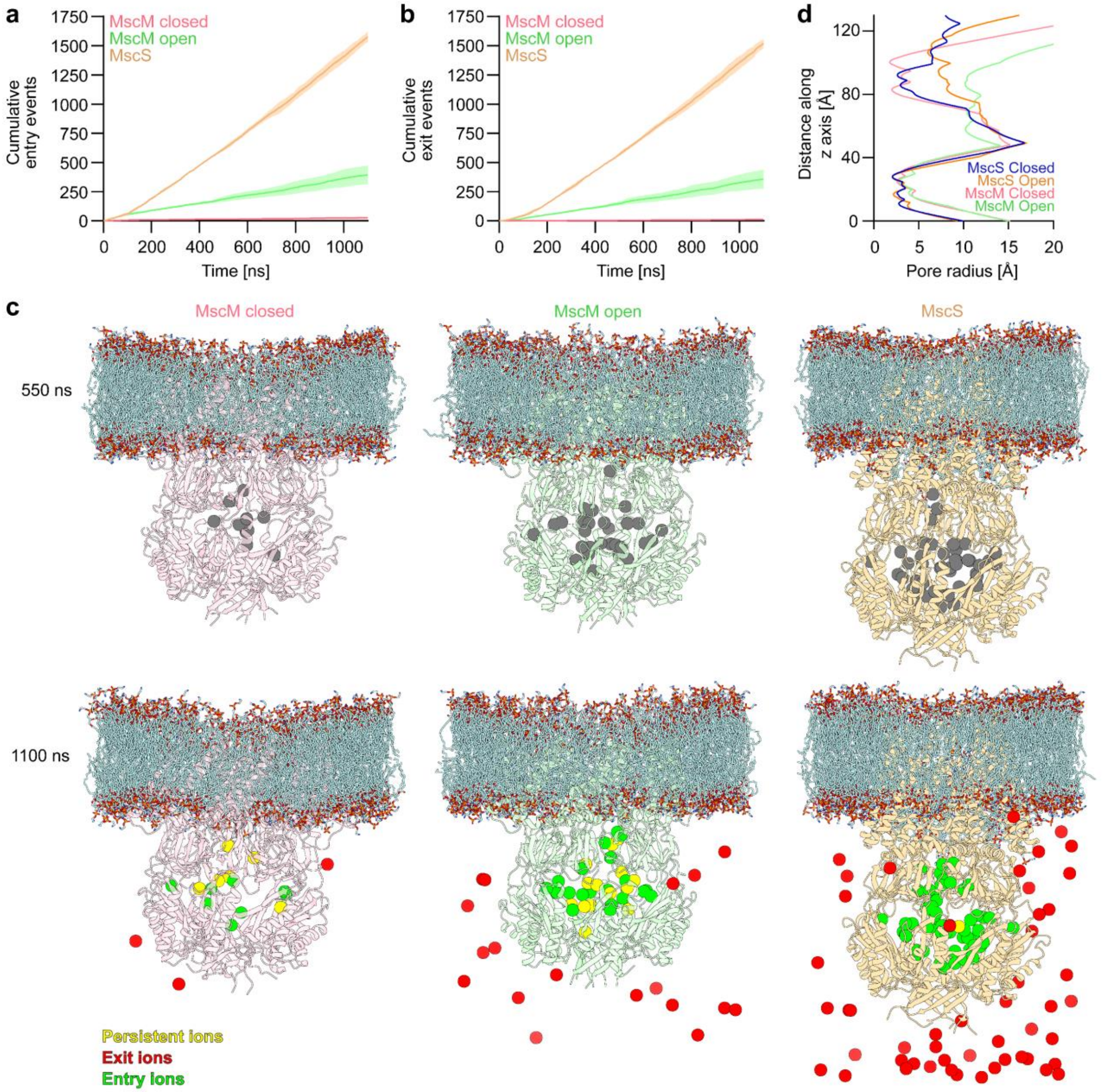

**Supplementary Figure 10. Analysis of the molecular-dynamics simulations. a,b** Plots of the cumulative entry (panel a) and exit (panel b) events through the fenestrations of MscS and MscM with closed and open fenestrations. **c** Snapshots from the mid-point (top panels) and end-point (bottom panels) of the simulations of MscM with closed (left panels) and open (middle panels) fenestrations and MscS (right panels). Top panels: Ions located in the cytoplasmic domains of the channels are shown as gray spheres. Bottom panels: Ions that were located in the cytoplasmic domains in the snapshots at the mid-point of the simulations and were still located in the cytoplasmic domains are shown as yellow spheres. Ions that were located in the cytoplasmic domains in the snapshots at the mid-point of the simulations and had exited the cytoplasmic domains are shown as red spheres. Ions that were not located in the cytoplasmic domains in the snapshots at the mid-point of the simulations and had entered the cytoplasmic domains are shown as green spheres. **d** Analysis of the pore radius of MscS and MscM in the closed and open conformations using HOLE[23].

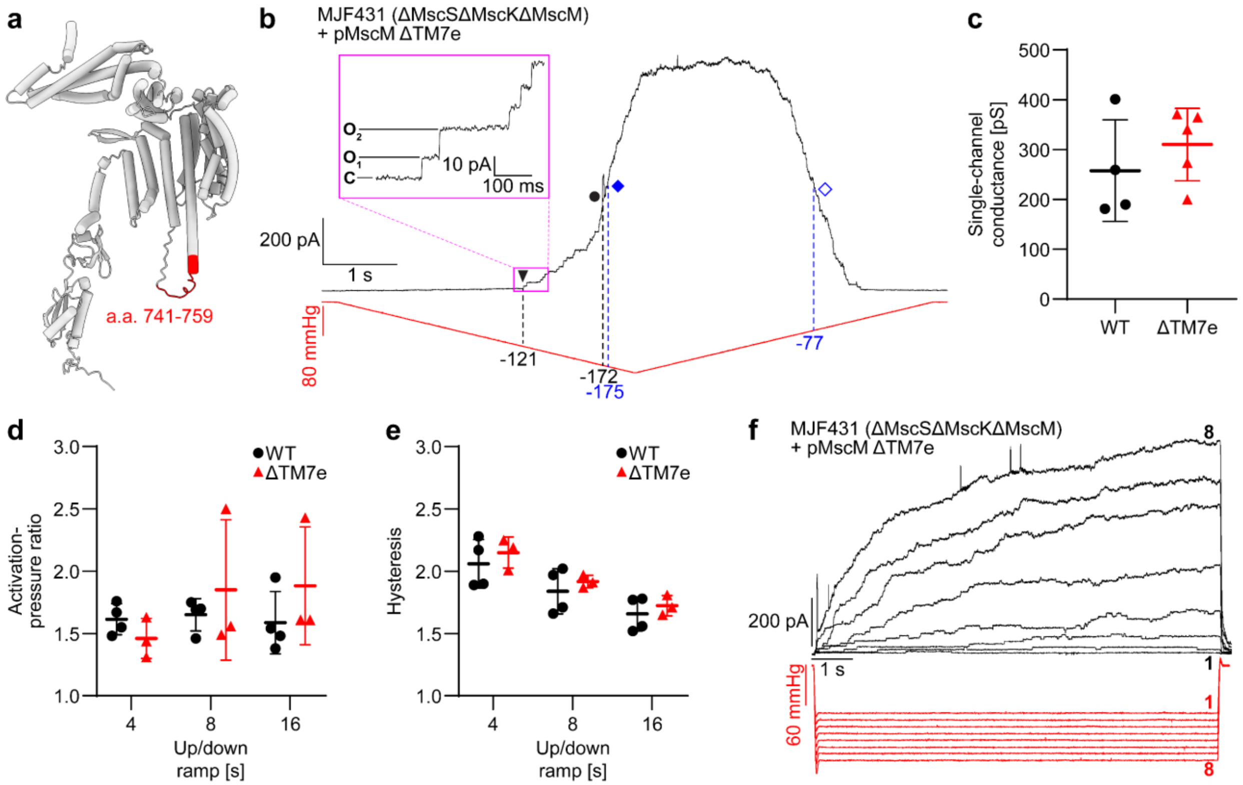

**Supplementary Figure 11. Electrophysiological characterization of the MscM ΔTM7e mutant. a** AlphaFold model of a single MscM subunit (white) indicating the region of the cytoplasmic extension of TM7 and the TM7-8 loop that was deleted (red) in the MscM ΔTM7e mutant. **b** Representative patch-clamp recording (n = 5) from a giant spheroplast generated from *E. coli* strain MJF431 (which does not express MscS, MscK and MscM) overexpressing MscM ΔTM7e in response to a symmetric 8-s-long pressure ramp from 0 mmHg to -210 mmHg and back to 0 mmHg. Currents were recorded in the inside-out excised patch configuration with a pipette potential of +30 mV. Pressure applied to the patch membrane and the recorded current are shown in red and black, respectively. Black triangle: first opening of MscM; black circle: first opening of MscL; filled blue rhombus: mid-point activation of MscM; empty blue rhombus: mid-point deactivation of MscM; dashed black lines: pressures at which the first MscM and the first MscL channels open; dashed blue lines: pressures at which the mid-point activation and mid-point deactivation of MscM occur. The inset shows a zoomed-in view of the MscM currents in the region indicated by the purple box. C: current level when all MscM channels are closed; $O_1$-$O_2$: current levels when one or two MscM channels are open. **c** Quantitative analysis shows no significant change in the single-channel conductance of MscM ΔTM7e (red triangles) compared to that of WT MscM (black circles). **d** Quantitative analysis shows no significant change in the activation-pressure ratio of MscM ΔTM7e (red triangles) compared to that of WT MscM (black circles) in response to ramps of different speeds. **e** Quantitative analysis shows no significant change in the hysteresis ratio of MscM ΔTM7e (red triangles) compared to that of WT MscM (black circles) in response to ramps of different speeds. **f** Representative patch-clamp recording (n = 5) from a giant spheroplast generated from *E. coli* strain MJF431 overexpressing MscM ΔTM7e in response to 10-s-long pulses with pressure increasing from -100 mmHg to -160 mmHg in -10 mmHg intervals. The ΔTM7e deletion does not alter the slow activation kinetics of MscM or its lack of desensitization/inactivation.

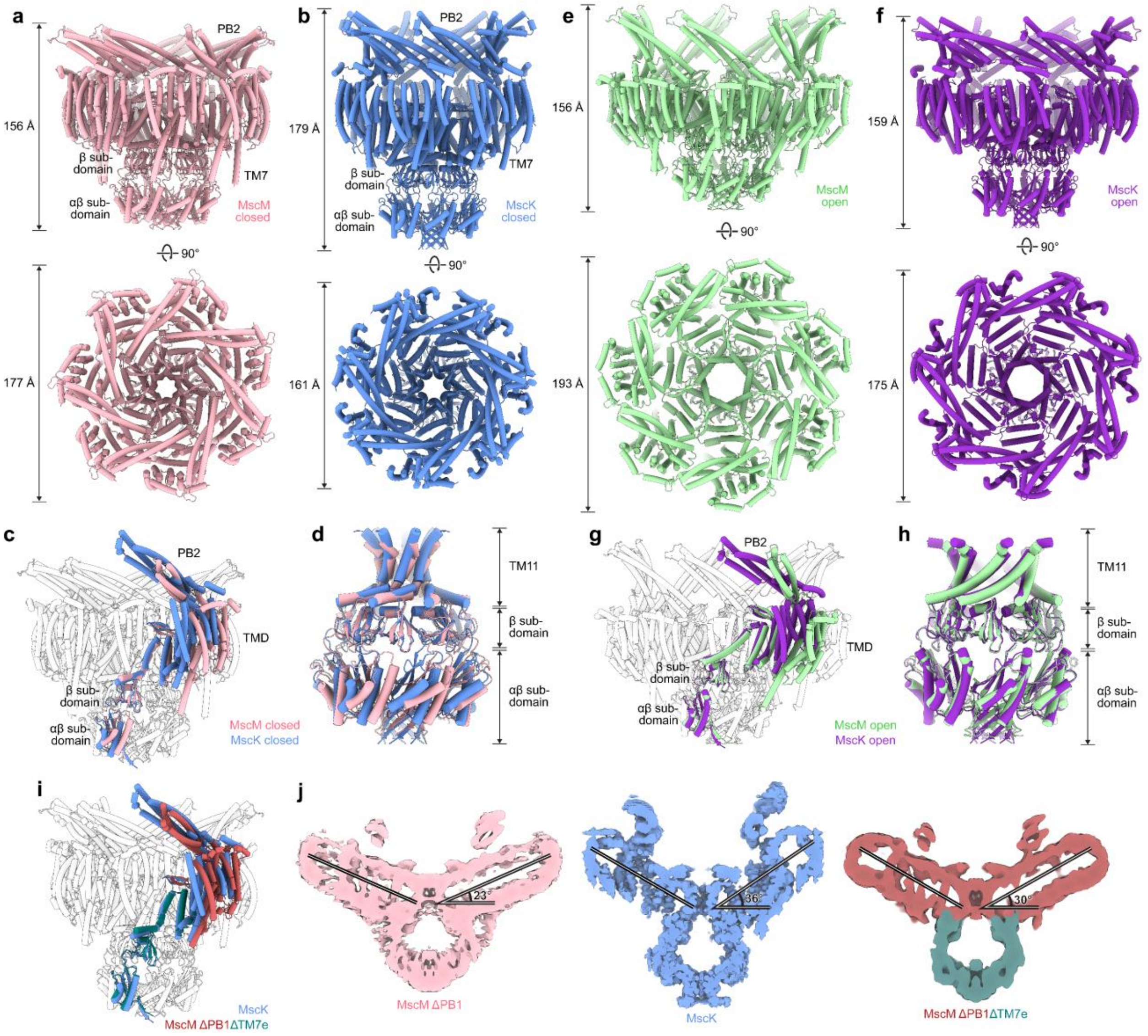

**Supplementary Figure 12. Structural comparison of MscM with MscK. a,b** Views parallel (top panels) and perpendicular (bottom panels) to the membrane of MscM (pink, left panels) and MscK[16] (PDB: 7uw5) (blue, right panels) in the closed conformation. **c** Alignment of the closed-conformation structures of MscM (one subunit in pink and six subunits in white) and MscK (one subunit in blue). **d** Alignment of TM11 and the cytoplasmic domain of MscM (pink) and MscK (blue) in the closed conformation based on TM11 shows that the two cytoplasmic domains are in different conformations. **e,f** Views parallel (top panels) and perpendicular (bottom panels) to the membrane of MscM (green, left panels) and MscK[16] (PDB: 7ux1) (purple, right panels). **g** Alignment of the open-conformation structures of MscM (one subunit in green and six subunits in white) and MscK (one subunit in purple). **h** Alignment of TM11 and the cytoplasmic domain of MscM (green) and MscK (purple) in the open conformation based on TM11 shows that the conformation of the cytoplasmic domain of MscM becomes the same as that of the cytoplasmic domain of MscK. **i** Alignment of the structures of MscM ΔPB1ΔTM7e (periplasmic and TM domains of one subunit in red and the cytoplasmic domain of the same subunit in teal, with the other six subunits in white) and MscK in the closed conformation (one subunit in blue) showing that the structure of MscM ΔPB1ΔTM7e more closely resembles that of MscK than that of MscM ΔPB1 (panel c). **j** Cut-away views parallel to the membrane of the cryo-EM maps of MscM ΔPB1 (pink, left panel), MscK[16] (EMDB: 26823) (blue, middle panel), and MscM ΔPB1ΔTM7e (red and teal, right panel) showing that truncation of the cytoplasmic extension of TM7 increases the curvature of the TM domain so that it more closely resembles that seen for MscK.

**Supplementary Table 1 | Cryo-EM data collection, refinement and validation statistics for all cryo-EM maps and models**

| | **WT MscM NaCl Closed conformation Global refinement** | **WT MscM NaCl Closed conformation Local refinement** | **MscM ΔPB1 NaCl Closed conformation Global refinement** | **MscM ΔPB1 NaCl Closed conformation Local refinement** | **WT MscM KCl Open conformation Global refinement** | **WT MscM KCl Open conformation Symmetry expansion** |
|---|---|---|---|---|---|---|
| **EMDB ID** | EMD-76005 | EMD-76007 | EMD-76009 | EMD-76010 | EMD-76011 | EMD-76013 |
| **PDB ID** | | 11SM | 11SO | | | |
| **Data collection** | | | | | | |
| **Microscope** | Titan Krios | Titan Krios | Titan Krios | Titan Krios | Titan Krios | Titan Krios |
| **Voltage (kV)** | 300 | 300 | 300 | 300 | 300 | 300 |
| **Camera** | K3 | K3 | K3 | K3 | K3 | K3 |
| **Magnification** | 105,000 | 105,000 | 105,000 | 105,000 | 105,000 | 105,000 |
| **Pixel size** | 0.847 | 0.847 | 0.847 | 0.847 | 0.847 | 0.847 |
| **Exposure time (s)** | 1.2 | 1.2 | 1.2 | 1.2 | 1.2 | 1.2 |
| **Frame rate** | 0.03 | 0.03 | 0.03 | 0.03 | 0.03 | 0.03 |
| **Electron exposure ($e^-$/Å$^2$)** | 46.21 | 46.21 | 46.84 | 46.84 | 46.84 | 46.84 |
| **Defocus range (μm)** | -0.8 to -2.0 | -0.8 to -2.0 | -0.8 to -2.0 | -0.8 to -2.0 | -0.8 to -2.0 | -0.8 to -2.0 |
| **Movie stacks (no.)** | 41,447 | 41,447 | 83,019 | 83,019 | 55,671 | 55,671 |
| **Reconstruction** | | | | | | |
| **Box size (pixels)** | 450 | 450 | 450 | 450 | 450 | 450 |
| **Initial particle images (no.)** | 7,016,041 | 7,016,041 | 11,459,225 | 11,459,225 | 5,857,401 | 5,857,401 |
| **Final particle images (no.)** | 10,378 | 50,627 | 58,387 | 61,183 | 136,448 | 135,809*** |
| **Symmetry imposed** | C7 | C7 | C7 | C7 | C1 | C1 |
| **Map resolution (Å)** | 6.7 | 3.4 | 4.3 | 3.4 | 2.8 | 3.2 |
| **FSC threshold** | 0.143 | 0.143 | 0.143 | 0.143 | 0.143 | 0.143 |
| **Map sharpening *B* factor (Å$^2$)** | Variable* | Variable* | Variable* | Variable* | Variable* | Variable* |
| **Map resolution range (Å)** | 4-8 | 3-7 | 3-7 | 3-6 | 2-6 | 2-6 |
| **Model composition** | | | | | | |
| **Non-hydrogen atoms** | | 10,255 | 31,598 | 10,325 | | 4,056 |
| **Protein residues** | | 1,309 | 5,628 | 1,323 | | 624 |
| **Ligands** | | 0 | 0 | 0 | | 0 |
| **Refinement** | | | | | | |
| **Initial model used** | | AlphaFold | AlphaFold | 11SO** | | 11SQ** |
| **Map-to-model CC (mask)** | | 0.78 | 0.64 | 0.80 | | 0.82 |
| **Map-to-model CC (volume)** | | 0.77 | 0.63 | 0.78 | | 0.81 |
| **Model-resolution (Å)** | | 3.9 | 7.6 | 3.7 | | 3.0 |
| **FSC threshold** | | 0.5 | 0.5 | 0.5 | | 0.5 |
| **Mean *B* factors (Å)** | | | | | | |
| **Protein** | | 118.72 | 296.33 | 123.49 | | 173.78 |
| **Ligand** | | N/A | N/A | N/A | | N/A |
| **R.m.s deviations** | | | | | | |
| **Bond lengths (Å)** | | 0.004 | 0.009 | 0.007 | | 0.004 |
| **Bond angles** | | 0.792 | 1.490 | 1.352 | | 0.740 |
| **Validation** | | | | | | |
| **MolProbability score** | | 1.57 | 1.91 | 1.90 | | 1.60 |
| **Clashscore** | | 11.40 | 17.01 | 18.54 | | 10.43 |
| **Ramachandran plot** | | | | | | |
| **Favored (%)** | | 98.38 | 96.99 | 97.33 | | 97.74 |
| **Allowed (%)** | | 1.54 | 1.38 | 1.07 | | 0.65 |
| **Outliers (%)** | | 0.08 | 1.63 | 1.60 | | 1.61 |
| **Rotamer outliers (%)** | | 0.81 | 0.63 | 0.63 | | 0.00 |
| **C-beta deviations (%)** | | 0.00 | 0.53 | 0.56 | | 0.00 |

| | WT MscM<br>KCl<br>Open<br>conformation<br>PB2 density map | WT MscM<br>KCl<br>Open<br>conformation<br>Composite map | MscM ΔPB1<br>KCl<br>Open<br>conformation<br>Local refinement | MscM ΔPD1ΔTM7e<br>NaCl<br>Closed<br>conformation<br>Global refinement | MscM ΔPD1ΔTM7e<br>NaCl<br>Closed<br>conformation<br>Local refinement |
|---|---|---|---|---|---|
| **EMDB ID** | EMD-76015 | EMD-76016 | EMD-76017 | EMD-76018 | EMD-76019 |
| **PDB ID** | | 11SQ | | 11SR | |
| **Data collection** | | | | | |
| **Microscope** | Titan Krios | Titan Krios | Titan Krios | Titan Krios | Titan Krios |
| **Voltage (kV)** | 300 | 300 | 300 | 300 | 300 |
| **Camera** | K3 | K3 | K3 | K3 | K3 |
| **Magnification** | 105,000 | 105,000 | 105,000 | 105,000 | 105,000 |
| **Pixel size** | 0.847 | 0.847 | 0.847 | 0.847 | 0.847 |
| **Exposure time (s)** | 1.2 | 1.2 | 1.2 | 1.2 | 1.2 |
| **Frame rate** | 0.03 | 0.03 | 0.03 | 0.03 | 0.03 |
| **Electron exposure ($e^-$/Å$^2$)** | 46.84 | 46.84 | 46.84 | 46.84 | 46.84 |
| **Defocus range (μm)** | -0.8 to -2.0 | -0.8 to -2.0 | -0.8 to -2.0 | -0.8 to -2.0 | -0.8 to -2.0 |
| **Movie stacks (no.)** | 55,671 | 55,671 | 11,623 | 26,496 | 26,496 |
| **Reconstruction** | | | | | |
| **Box size (pixels)** | 450 | 450 | 450 | **450** | **450** |
| **Initial particle images (no.)** | 5,857,401 | | 1,298,131 | 4,442,810 | 4,442,810 |
| **Final particle images (no.)** | 69,165 | | 34,087 | 48,883 | 57,877 |
| **Symmetry imposed** | C7 | C7 | C7 | C7 | C7 |
| **Map resolution (Å)** | 3.3 | 3.2**** | 3.4 | 5.1 | 3.5 |
| **FSC threshold** | 0.143 | 0.143 | 0.143 | 0.143 | 0.143 |
| **Map sharpening *B* factor (Å$^2$)** | 0 | Variable* | Variable* | Variable* | Variable* |
| **Map resolution range (Å)** | 3-8 | 2-6 | 3-5 | 4-8 | 3-5 |
| **Model composition** | | | | | |
| **Non-hydrogen atoms** | | 28,392 | | 32347 | 13650 |
| **Protein residues** | | 4,368 | | 5537 | 1771 |
| **Ligands** | | 0 | | 0 | 0 |
| **Refinement** | | | | | |
| **Initial model used** | | This study | | AlphaFold | 11SR** |
| **Map-to-model CC (mask)** | | 0.84 | | 0.55 | 0.76 |
| **Map-to-model CC (volume)** | | 0.83 | | 0.53 | 0.74 |
| **Model-resolution (Å)** | | 3.4 | | 8.3 | 3.94 |
| **FSC threshold** | | 0.5 | | 0.5 | 0.5 |
| **Mean *B* factors (Å)** | | | | | |
| **Protein** | | 173.28 | | 536.24 | 125.35 |
| **Ligand** | | N/A | | N/A | N/A |
| **R.m.s deviations** | | | | | |
| **Bond lengths (Å)** | | 0.004 | | 0.004 | 0.009 |
| **Bond angles** | | 0.742 | | 0.897 | 1.049 |
| **Validation** | | | | | |
| **MolProbability score** | | 1.62 | | 1.96 | 1.84 |
| **Clashscore** | | 11.08 | | 11.77 | 12.25 |
| **Ramachandran plot** | | | | | |
| **Favored (%)** | | 97.74 | | 94.52 | 96.39 |
| **Allowed (%)** | | 0.65 | | 4.71 | 2.81 |
| **Outliers (%)** | | 1.61 | | 0.76 | 0.80 |
| **Rotamer outliers (%)** | | 0.00 | | 0.62 | 0.69 |
| **C-beta deviations (%)** | | 0.00 | | 0.00 | 0.00 |

* Map was sharpened with EMReady
** Refinement and validation statistics were reported using only the part of the structure built using the corresponding cryo-EM, as discussed in the Methods
*** After symmetry expansion
**** Accurate Gold-standard FSC resolution could not be assigned for composite map

**SUPPLEMENTARY MOVIES**

**Supplementary Movie 1. Movie highlighting the different domains of MscM followed by a morph between MscM in the closed and open conformations viewed parallel (left) and perpendicular (right) to the membrane, illustrating the conformational changes associated with channel gating.**

**Supplementary Movie 2. Movie showing a morph between the MscS-like core of MscM in the closed and open conformations, illustrating the conformational change of the cytoplasmic domain and the opening of the fenestrations.**

**Supplementary Movie 3. Movie showing a morph between MscM with a full-length TM7 and ΔTM7e viewed parallel (left) and perpendicular (right) to the membrane, illustrating the role of TM7e in coupling the conformations of the transmembrane and cytoplasmic domains.**

**Supplementary Movie 4. Movie showing morphs for MscM (left) and MscK (right) between their closed and open conformations, illustrating the conformational changes associated with gating of the two channels.**
